# Thermogenetics for cardiac pacing

**DOI:** 10.1101/2024.01.02.573885

**Authors:** Alexander V. Balatskiy, Alexey M. Nesterenko, Vera S. Ovechkina, Aleksandr A. Lanin, David Jappy, Rostislav A. Sokolov, Vitalii D. Dzhabrailov, Elena A. Turchaninova, Mihail M. Slotvitsky, Elena S. Fetisova, Alexander A. Moshchenko, Sofya K. Andrianova, Ruslan M. Karpov, Semyon S. Sabinin, Diana Z. Biglova, Andrey V. Zakharov, Georgy M. Solius, Ekaterina M. Solyus, Sergei V. Korolev, Oleg V. Podgorny, Valeriya A. Tsvelaya, Konstantin I. Agladze, Ilya V. Kelmanson, Andrei B. Fedotov, Andrei V. Rozov, Tobias Bruegmann, Alexei M. Zheltikov, Andrey A. Mozhaev, Vsevolod V. Belousov

## Abstract

Cardiac arrhythmias are common disorders that can be fatal. Modern methods of treating bradyarrhythmias include the implantation of pacemakers and cardioverter–defibrillators. However, implantable devices can cause various complications related to the electrodes installed inside the heart, including infection. Less invasive heart rhythm modulation could be beneficial for some cohorts of patients. Here, we demonstrate an alternative approach to heart pacing based on thermogenetics. We used adeno-associated viruses to deliver genetic human transient receptor potential subfamily V member 1 (TRPV1), a heat-sensitive cation channel, into isolated cardiomyocytes and the mouse heart. This allowed us to induce action potentials and control contractility using short heat pulses delivered by infrared laser illumination. Using this approach, we demonstrated the thermogenetic pacing of isolated cardiomyocytes *in vitro* and in the mouse heart *in vivo*. Our results demonstrate the potential of thermogenetics for developing therapeutic strategies for heart rhythm modulation.

---

Electrical stimulation of cardiomyocytes and cardiac tissue is the gold standard for laboratory studies and clinical applications. In patients, implantation of electrical devices, such as pacemakers or cardioverter-defibrillators, is very common, in some countries the implantation rate is >1000 cases per million people^1^. According to some estimates, more than 1 million devices are implanted worldwide annually^2^. Nevertheless, electronic pacemakers have limitations, such as the need for an invasive surgical procedure, the risk of infection, device failure, regular battery replacement, and electrochemical reactions at the electrode contact sites. In addition, there is the possibility of endocarditis, thrombosis, electrode dislocation, and other complications both in the early postoperative and long-term periods. Automatic cardioverter-defibrillators, when triggered, generate extremely painful discharges for the patient, leading to decreased quality of life. In a recent study, major complications occurred in 8.2% of patients after the implantation of a pacemaker or defibrillator within 90 days of hospital discharge^3^.

Thus, there is a need for alternative approaches to reduce the complications associated with device implantation. Approaches using gene and cell therapy might be used as alternative solutions for the correction of cardiac arrhythmias^4-6^. The emergence of a variety of genetically encoded tools has opened the possibility of studying and controlling the activity of nerve, muscle, and secretory cells by exposing ectopically expressed molecular targets to chemical ligands, light, or temperature fluctuations. Technologies using such principles of action are known as chemogenetics, optogenetics, and thermogenetics, respectively. The advantage of these technologies is the possibility of targeting certain groups of cells with high spatial resolution using tissue-specific promoters.

Optogenetic methods using photosensitive proteins to control cardiomyocyte contractions have been applied to isolated cardiomyocytes^7,8^, zebrafish hearts^9^, transgenic mouse hearts^10^, and *ex vivo* mouse hearts with intramyocardial transgene delivery by adeno-associated viruses (AAV)^11,12^. Despite the existing options for the use of optogenetics on cardiomyocytes, this technology has some limitations and disadvantages. First, optogenetics uses rhodopsin ion channels that need to be activated by visible light. Research aimed at developing redshifted channelrhodopsins has not yet yielded effective deep-penetrating activation radiation since most studies still use visible light to activate the widely used channelrhodopsin 2 (ChR2)^13-15^. This leads to the fact that the stimulation of cells in optically opaque animals in the vast majority of cases is invasive, using implantable devices such as optical fibres. Despite approaches to the creation of rhodopsins with high conductivity^16^, most have low conductivity^17^ and require a high level of channel expression and a high intensity of activating light, which can lead to phototoxic effects. Second, a disadvantage that limits the use of the *in vivo* optogenetic approach is the use of channelrhodopsins which are not found in mammalian organisms and can cause an immune response and, as a result, rapid death of the cells expressing them^18,19^.

Thermogenetics is a promising alternative approach based on the use of non-selective cation channels, which can be opened by temperature changes^20^. The advantage of using these channels is that they are present in mammals, which makes an immune response much less likely. These channels are found in many tissues and cells, some are present on the membranes of sensory neurons, acting as thermal sensors, and in some cases, pain receptors, in various groups of animals. These channels also have a significantly higher conductance compared to ChR^21^. The high conductivity and ability of some to respond to small temperature shifts of 1–2°C have contributed to the use of these channels as thermogenetic activators of various cells. Another advantage of the thermogenetic approach is the wide range of ways to activate thermosensitive channels: infrared (IR) radiation^22-24^, microwaves^25^, focused ultrasound^26^ or magnetic nanoparticles in an alternating magnetic field^27^ can be used. Previously, the possibility of IR laser stimulation of single cells with the possibility of generating successive action potentials on a millisecond scale, commensurate with the response time of the light-sensitive channels used for optogenetics, was demonstrated^23^.

In the present study, we applied a thermogenetic approach to control the rhythm of isolated cardiomyocytes *in vitro*, and the mouse heart *in vivo* using pulsed IR laser heating, using human transient receptor potential subfamily V member 1 (hTRPV1) **(Fig. 1A)**. Our study demonstrates that TRPV1 is an excellent tool for thermogenetics and has the potential for a wide range of future applications from research to therapeutic interventions.

**Fig. 1.**
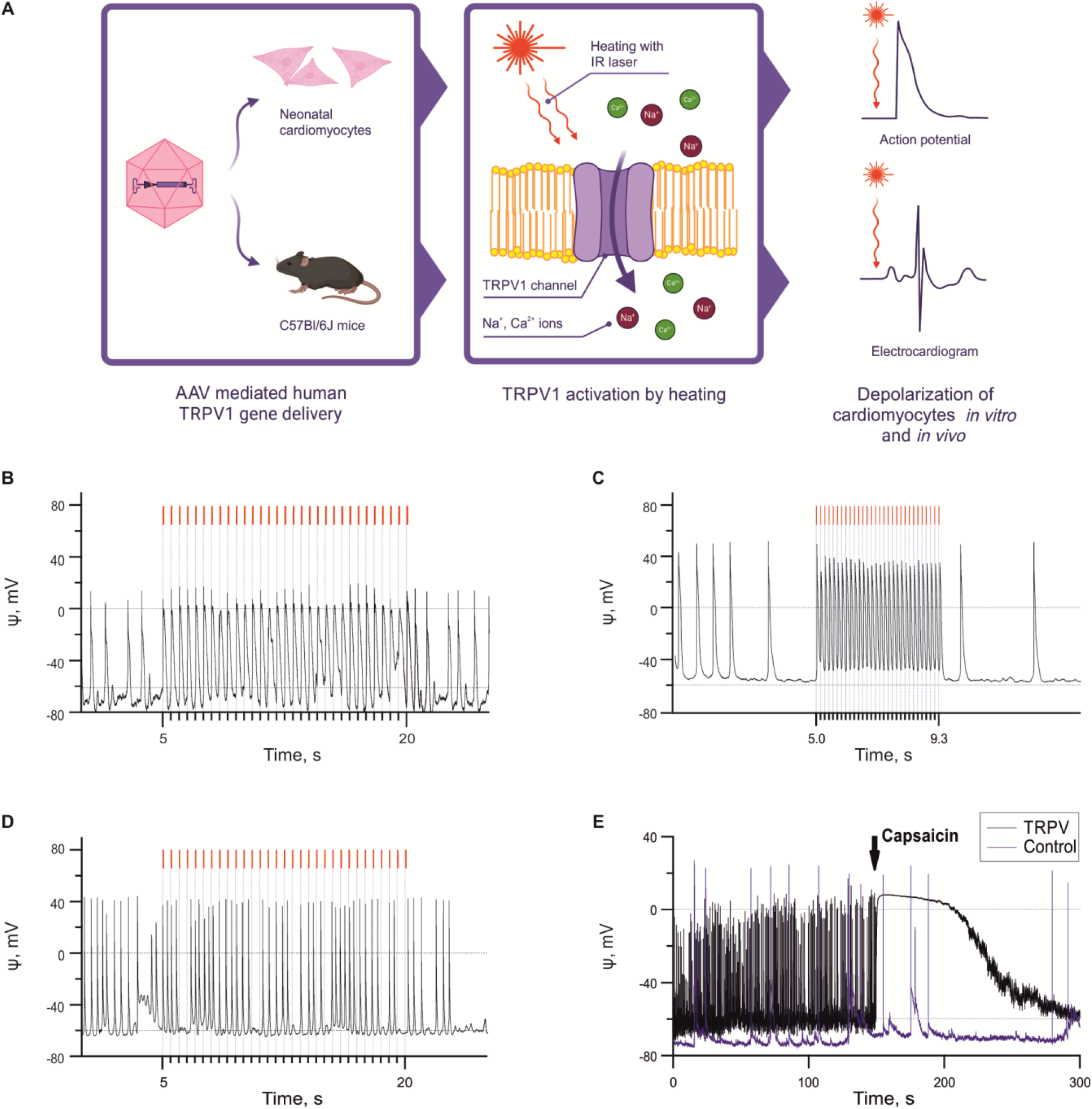
Experimental design and thermogenetic pacing of isolated neonatal murine cardiomyocytes using IR laser. **a**, Schematic overview of the thermogenetic experiments. **b**, Depolarization of cardiomyocytes transduced with hTRPV1(−109aa) induced heat pulses with a repetition rate of 2 Hz. **c**, Depolarization of cardiomyocytes expressing hTRPV1 induced by 7 Hz heat pulses. **d**, Heating of control cells transduced with GCaMP6s. In **b-d** the laser pulses are shown in red. **e**, Addition of capsaicin to cells transduced with hTRPV1, and to control cells transduced with GCaMP6s.

## Results

We demonstrated the possibility of thermogenetic pacing in vitro on isolated murine neonatal cardiomyocytes and in vivo in adult mice.

### Thermogenetic pacing of cardiomyocytes in vitro

We chose the hTRPV1 channel as a thermogenetic instrument. When heated to 41–43°C, it opens and passes Na^+^ and Ca^2+^ through the plasma membrane into the cell, which leads to depolarization. We hypothesized that this depolarization can trigger action potentials in cardiomyocytes. To demonstrate thermogenetic activation *in vitro*, we expressed the truncated version of hTRPV1 in neonatal murine cardiomyocytes. Since the fill-size hTRPV1 is too big to pack into AAV with a fluorescent protein, we transduced cells with AAV-PHP.S containing cTnT_hTRPV1(–109aa)_P2A_mRuby. This truncated channel demonstrated the same activity as the full-length hTRPV1. The activation of hTRPV1(−109aa) in HEK293 cells heated by a nichrome wire loop is shown in **Supplementary Fig. 1**.

To heat cardiomyocytes, we used an IR laser connected to an optical fiber fixed to a micromanipulator. We gradually increased the pulse duration and power until pacing was achieved. The average pulse characteristics used for successful pacing are given in **Supplementary Table 1** and are shown in detail in **Supplementary Fig. 2**. The entire dataset is deposited online^28^. Electrophysiological measurements demonstrated action potentials (AP) induced by laser-induced heat pulses at frequencies up to 7 Hz **(Fig. 1b, c)**. This coupling was absent in control cells (non-transduced or transduced with cTnT_GCaMP6s, **Fig. 1d**). Application of 100 µM capsaicin through a pipette within about 20 μm of the recorded cell led to rapid and long-lasting depolarization in cells transduced with cTnT_hTRPV1(−109aa)_P2A_mRuby but not in the control cells **(Fig. 1e)**. All inward currents were completely abolished by the selective TRPV1 inhibitor AMG517 (1 µM solution applied through a pipette near the cell), **(Supplementary Fig. 3)**. The pulse width and the laser power do not determine the outcome of low-frequency (2 Hz) pacing: for the two-factor logit model, LLR p-value = 0.5. The success ratio for high-frequency (7 Hz) pacing grows significantly with laser power (p-value < 0.05) and decreases with pulse width with a lower significance (0.05 < p-value < 0.1); for the two-factor logit model LLR p-value = 0.006.

To assess the effect of TRPV1 activation on the recruitment of various membrane conductances involved in the generation of AP, we analyzed the kinetics of spontaneous APs and APs evoked by activation of hTRPV1(−109aa) channels **(Supplementary Fig. 4)**. Although the amplitude, depolarization rate, half-width, and repolarization rate were slightly but significantly different in spontaneous and TRPV1-evoked APs, the AP waveforms were similar in both groups, suggesting that the generation of thermogenetically evoked APs involves the same types of voltage-gated channels and the same sequence of activation as natural APs.

To confirm our ability to stimulate cardiac cells in vitro, we performed optical mapping on a monolayer of murine neonatal cardiomyocytes using a Fluo-4 AM calcium indicator **(Fig. 2)**. Upon pulse heating with an IR laser, hTRPV1-FLAG-expressing cells were stimulated at a given frequency. This effect was not observed in the culture that did not express TRPV1. The wave propagation velocity did not differ between hTRPV1-FLAG-expressing and control monolayers **(Fig. 2g)**.

**Fig. 2.**
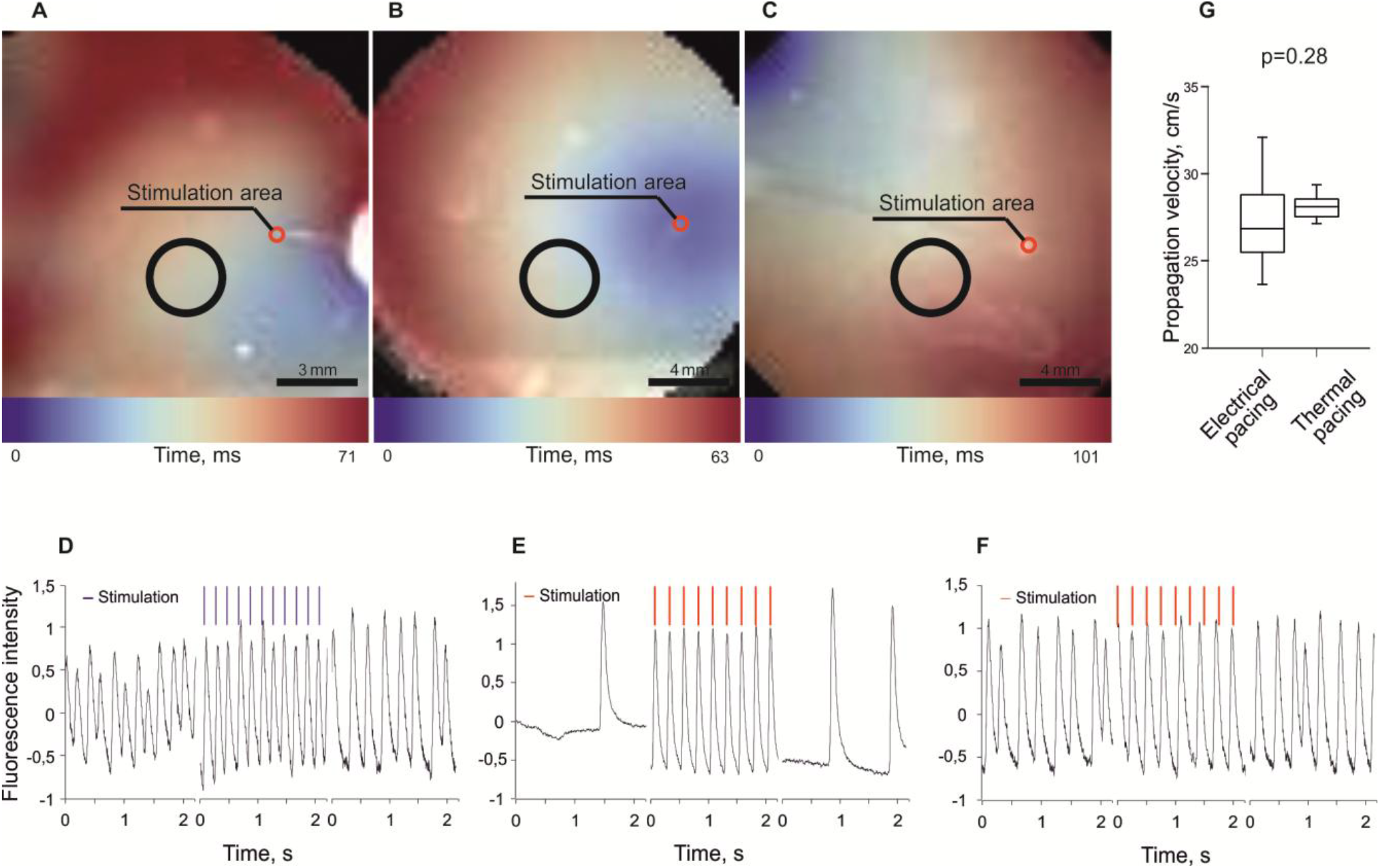
Optical mapping analysis of neonatal mouse cardiomyocyte monolayers. **a-c**. Activation maps of excitation wave propagation through the cardiomyocyte monolayer. Black circles show the regions where fluorescence intensity was measured. **d-f**. Plots showing Fluo-4 AM fluorescence intensity in specific regions of the monolayer. **a**,**d**, Excitation wave propagation across a monolayer of hTRPV1-FLAG-expressing cardiomyocytes, upon stimulation with a platinum electrode. **b**,**e**. Excitation wave propagation across a monolayer of hTRPV1-FLAG-expressing cardiomyocytes upon IR laser pulse heating. **c**,**f**. Excitation wave propagation across a monolayer of control cardiomyocytes upon IR laser pulse heating. **g**. Propagation velocities upon electrical stimulation and pulse heating.

### Thermogenetic heart pacing in vivo and ex vivo

To reliably detect hTRPV1 channel in mouse tissues, we decided to use AAVs carrying cTnT_hTRPV1-3×FLAG in the in vivo experiments. We first tested the functional activity of hTRPV1 fused with the FLAG peptide. We transfected HEK cells with pCAG_hTRPV1_p2A_GCaMP6s and pCAG_hTRPV1-3×FLAG_p2A_GCaMP6s plasmids and recorded inward currents at different temperatures **(Supplementary Fig. 5)** for both channels. hTRPV1-FLAG demonstrated smaller inward currents compared to native hTRPV1. However, unlike the native hTRPV1, for which we sometimes detected very small currents at 37°C, we did not detect any inward current at 37°C for hTRPV1-FLAG. These results suggest higher safety of the hTRPV1-FLAG channel.

All pacing experiments were performed four weeks after the injection of AAVs carrying cTnT_hTRPV1-3×FLAG. For in vivo experiments, 4 hTRPV1-FLAG-expressing mice and 3 control mice were used. Mice did not demonstrate signs of pathology up to 14 months after the injection. hTRPV1-FLAG channels were more or less evenly distributed in the myocardium **(Supplementary Fig. 6a)**, 90.7±1.8% of cardiomyocytes were infected in apexes, 76.0±2.6% were infected in other parts of the ventricular myocardium, and 79.6±5.3% were infected in atria. The percentage of the infected cells in heart apex was significantly higher than in other zones (p<0.001). We also discovered hTRPV1-FLAG expression in sinoatrial node cells stained with anti-HCN4 antibodies **(Supplementary Fig. 6g)**. Anti-FLAG staining was detected not only on the plasma membrane but also inside the cardiomyocytes **(Supplementary Fig. 6h)**. We did not find any off-target expression in liver, brain, lungs and skeletal muscles **(Supplementary Fig. 6c-f)**.

Open hearts of anesthetized and ventilated mice were heated with laser pulses. In animals transduced with cTnT_hTRPV1-3×FLAG, the shape of ventricular complexes changed (QRS and QTc durations increased, **Supplementary Fig. 7)** as was expected due to the transition from the sinus to the ventricular rhythm. Heartbeats were synchronized with heat pulses at frequencies up to 6 Hz **(Fig. 3a, b)**. In control mice, complex shapes were also slightly affected but the heart rate was not modified during IR laser treatment **(Fig. 3c, d)**. Note, synchronization (phase locking, “L” in **Fig. 3a, b**) occurred 10–20 sec after the beginning of pacing. In contrast to cell culture in a Petri dish, the heart is an object with a large thermal capacity, leading to a baseline temperature increase of up to 40°C at about 12 sec after the beginning of the laser pulse train irradiation. Above this drift, there are sharp temperature spikes with a magnitude of 3°C and a duration of 50 ms, the same as the laser pulse width. These small pulses allow hTRPV1 activation temperature threshold to be reached **(Fig. 3c–d)**. We used the same IR laser setup for experimental animals. With our setup, we had to use the maximum possible laser power to reach the target temperature in a short time, so we did not have much flexibility in choosing the heating parameters. The statistical analysis did not reveal any correlation between pulse characteristics or the initial frequency and successful pacing **(Supplementary Table 2)**.

**Fig. 3.**
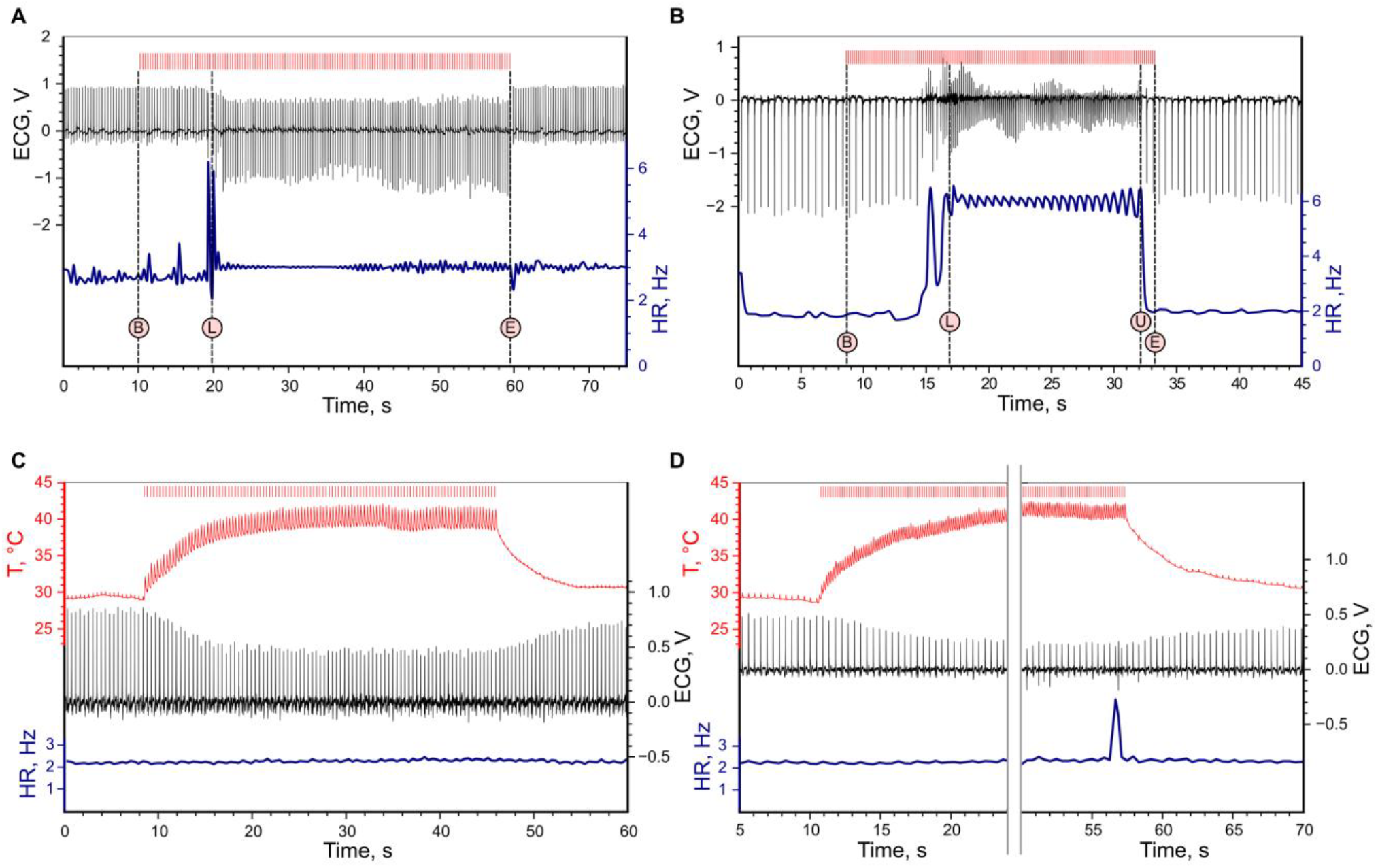
Thermogenetic heart pacing *in vivo* using apex heating by IR laser pulses. Denoised ECG (black lines) and heart rate (HR, blue lines) are depicted for experimental animals with hTRPV1 expression (**a**,**b**) and control animals (**c**,**d**). IR 3 Hz (**a**,**c**) and 6 Hz (**b**,**d**) laser pulses were applied (top red dashes). Temperature was measured only in control animals (red curve). Events are labelled with circles: ‘B’/’E’ — beginning and ending of pulse sequence, ‘L’/’U’ — phase locking/unlocking.

During pacing of the atrium, the imposed frequency was held very precisely and the phase locking was set very quickly **(Fig. 4)**. The QRS shape and ECG intervals stayed unchanged **(Supplementary Fig. 7)**. In the beginning, the P-wave arose quickly, even before the end of the laser pulse, so the laser pulse was close to the end of the R-R interval. **(Fig. 4c)**. The further the pacing progressed, the more the P-wave moved to the end of the pulse. Finally, the P-wave reached the position when its beginning corresponded approximately to the end of the laser pulse (**Fig. 4d)**. The described behavior is general and not unique to this sample, we provide different data samples deposited at ref. ^29^ (see also **Supplementary Table 2**). Initial phase locking probably occurred when the cells in the atrium reached a target temperature and a critical amount of cells depolarized. In this case, the P-peak was located approximately 40 ms after the laser pulse. After the action potential has already developed and the atrium has already contracted, prolonged heating can only cause a calcium overload. The heart might be able to adapt its behavior to develop action potentials only when the heating is already finished.

**Fig. 4.**
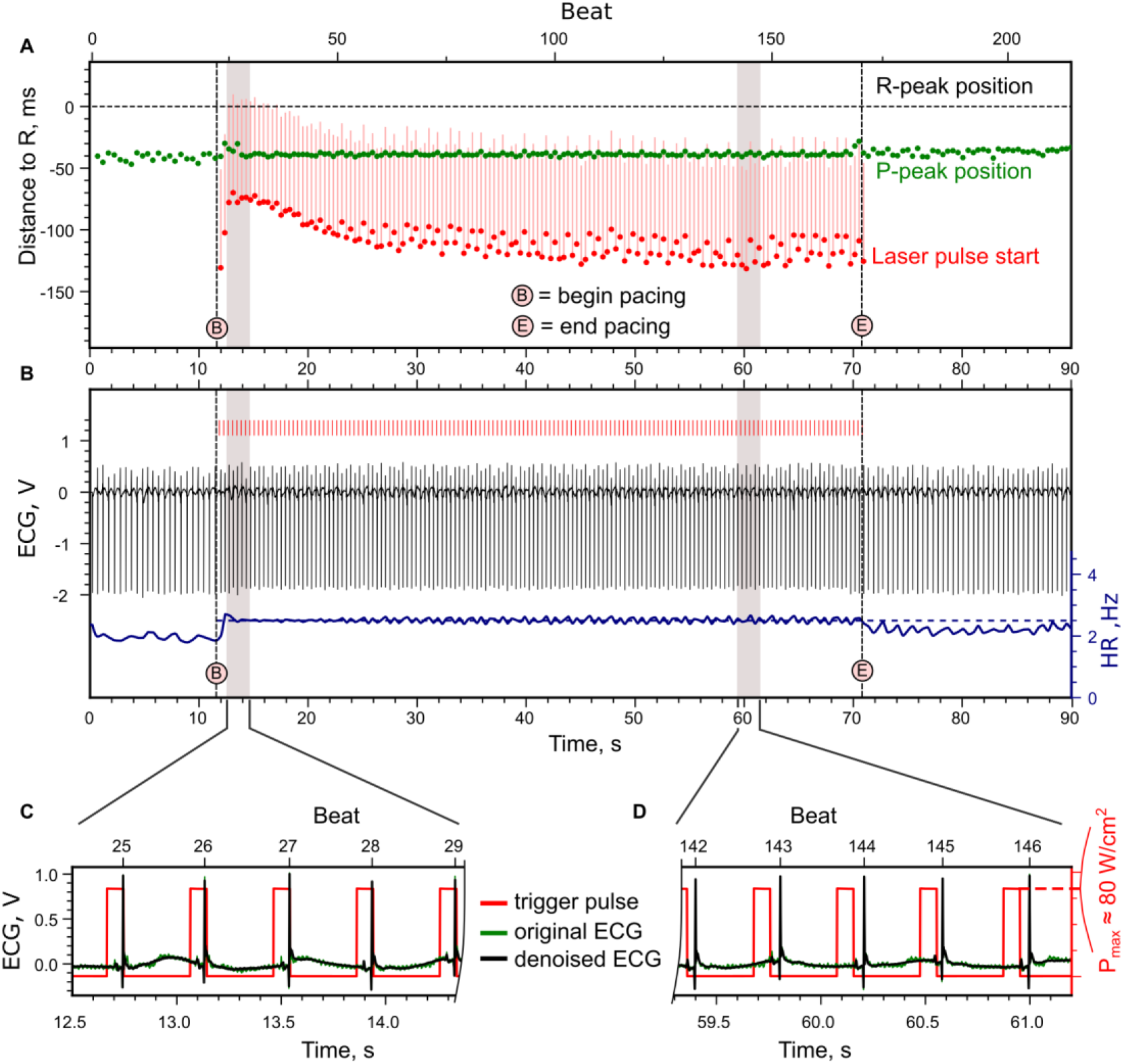
Typical example of atrium pacing with a frequency slightly higher than the intrinsic one. Pacing was at 2.5 Hz with 80 ms pulses, initial HR was ∼2 Hz. **a**, Position of the P-peak and laser pulse in relation to the R-peak in ms. Negative values mean that events occur *before* the reference peak. **b**, Change of complex shape (black curve) and heart rate (blue curve) during the experiment. Vertical red lines demonstrate the pulse widths of the trigger pulses. **c**,**d** ECG signal and trigger pulses in short intervals corresponding to the beginning and ending parts of the pacing. The positions of the intervals are indicated with grey bars in panes **a** and **b**.

In contrast to pulsed IR illumination, constant heating of the atrium by the IR laser induced a very unstable beat with a high variability in complex shape **(Supplementary Fig. 8c–c’)**. In addition, increased heart rate was also observed in control hearts as well. The reaction of control hearts during pacing with typical parameters is shown **in Supplementary Fig. 9**. Heart rate was considerably increased **(Supplementary Fig. 9b)**, however no synchronization of laser peaks with heart beating was detected **(Supplementary Fig. 9a**). Heating of ventricular tissues in control hearts did not affect the rhythm in both pulsed and continuous modes **Fig. 3c,d and Supplementary Fig. 10)**, but affected the amplitude of the R-peak (**Fig. 3c,d)**.

In control recordings, we were able to observe typical heating of cardiac tissues measured with a probe. In the case of atrium pacing, the phase locking in experimental animals was achieved in a much shorter time (in 1–2 seconds) than it took for the atrium to reach the threshold temperature (∼10 sec) **(Supplementary Fig. 9b)**. Cells at the surface of the atrium were probably heated quicker than the entire atrium, providing initial depolarization sufficient for pacing.

To better validate our method and measure conduction velocities, we performed optical mapping experiments on isolated mouse hearts. For these experiments, five mice injected with cTnT_hTRPV1-3×FLAG, and five control animals were used. We successfully stimulated different areas of the heart, including both atria and both ventricles. A representative example of ventricular pacing is shown in **Fig. 5**.

**Fig. 5.**
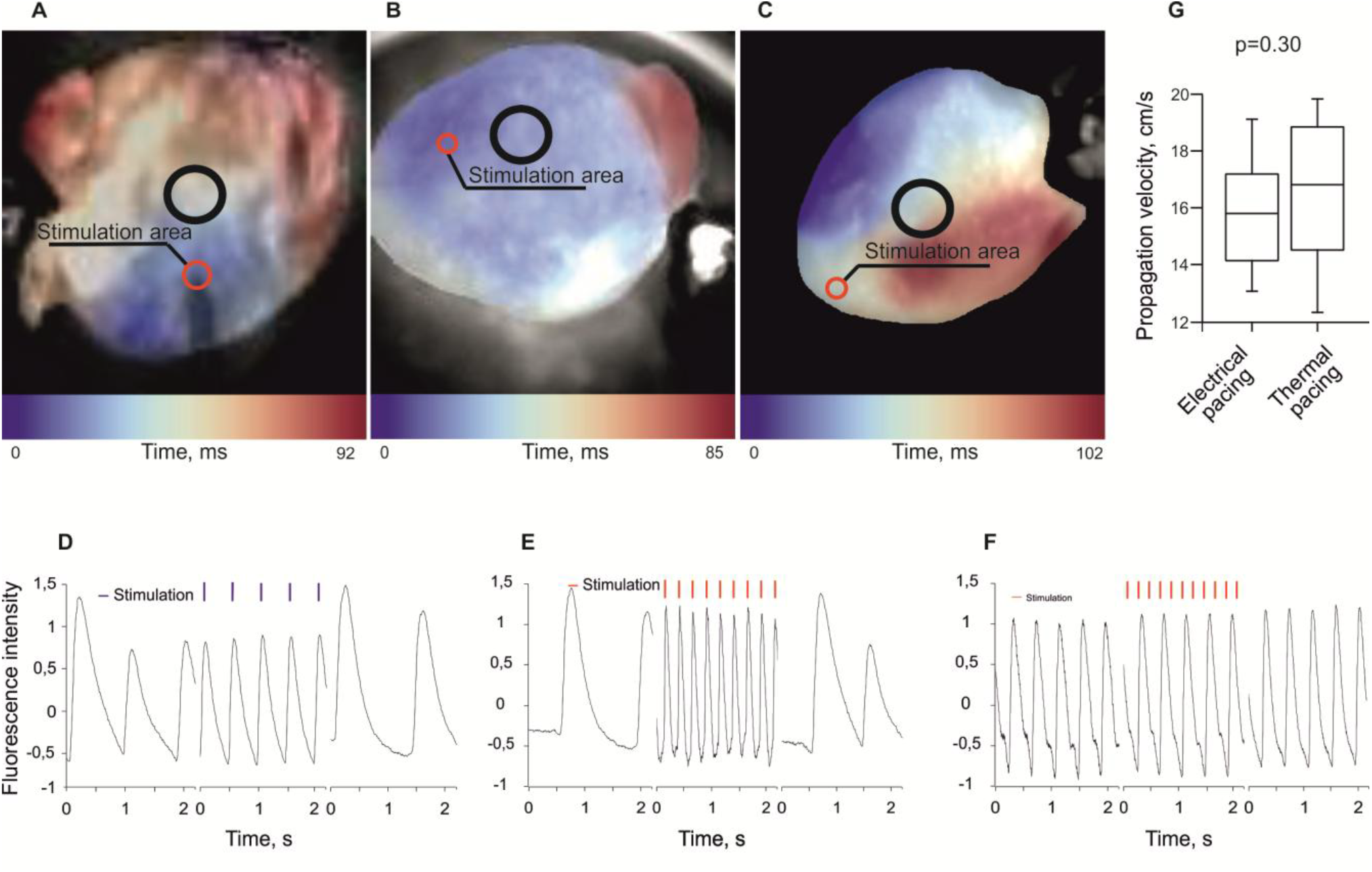
Optical mapping analysis of mouse hearts. **a-c**. Activation maps of excitation wave propagation in the hearts. Black circles show the regions where fluorescence intensity was measured. **d-f**. Plots showing Fluo-4 AM fluorescence intensity in specific regions of the hearts. **a**,**d**, Excitation wave propagation in hTRPV1-FLAG-expressing hearts upon stimulation with a platinum electrode. **b**,**e**. Excitation wave propagation in hTRPV1-FLAG-expressing hearts upon IR laser pulse heating. **c**,**f**. Excitation wave propagation in control hearts upon IR laser pulse heating. **g**. Propagation velocities upon electrical stimulation and pulse heating.

We achieved quite a high spatial resolution, making it possible to induce pacing with a 1.8-2.2 mm heating spot applied to the desired area **(Supplementary Fig. 12)**. In these experiments, we also aimed to determine the optimal stimulation parameters. Due to the technical limitations of our setup, pacing was only possible with the maximum laser power allowed by the laser driver. Lower power values required too long pulses to reach the necessary temperature. At the given power of approximately 69 W/cm^2^, we determined the minimal duty cycle required for successful pacing – 35%, which corresponds to 87.5 ms for stimulation at 4 Hz.

### Rhythm imposing peculiarities and calcium dynamics in cardiomyocytes

For most patients, heart pacing therapy is required chronically. Therefore, it is important to evaluate potential problems of a new technology that should be addressed before it is transferred to the clinic. For thermogenetic heart pacing, we report two major problems observed so far: incomplete phase locking and loss of phase locking during long-time pacing.

We recorded a lot of samples with accurate heart pacing **(Supplementary Table 2)** with S:R = 1:1 phase locking (S : R means Stimulus : Response). However, in many cases phase locking was incomplete, some laser pulses initiated a heart beat and others were skipped. In atrium pacing experiments, there were events of incomplete phase locking with S:R being 4:3 or 3:2 **(Supplementary Fig. 13)**.

During the pacing series longer than 40 s, especially when the ventricle was paced, we observed spontaneous loss of synchronization. In some cases, pacing was spontaneously restored **(Fig. 6)**. This phenomenon was not observed during conventional electric pacing **(Supplementary Fig. 14)**. One of the possible causes of pacing loss is overheating, because the heart may not cool effectively in an open chest. However, spontaneous pacing restoration during heating does not support this hypothesis. Long pacing of isolated cardiomyocytes demonstrated a similar effect: the pacing was spontaneously lost and restored during a long series, but finally, cells stopped responding **(Fig. 7a)**. Since TRPV1 has significant conductivity for Ca^2+^ ions, we hypothesized that the possible cause of response termination was calcium overload. Immunostaining of tagged hTRPV1 demonstrated a significant fraction is localized in the endoplasmic reticulum (ER). ER and tagged hTRPV1 double staining with colocalization analysis gave us tresholded Mandel’s coefficients tM1 = 0.292 and tM2 = 0.754, which means that almost 30% of all hTRPV1 is localized in ER, and 75% of the entire ER contains hTRPV1 **(Fig. 7b–e)**. We used the genetically encoded calcium sensor GCaMP6s to visualize calcium dynamics inside cardiomyocytes both in calcium-containing and calcium-free media. In both cases, after the initial synchronization of calcium spikes with heat pulses, cells then lost synchronization, and demonstrated a rise in intracellular calcium together with loss of calcium spiking activity **(Fig. 7f)**. This data demonstrates that ER-localized hTRPV1 may allow calcium to move from the ER to the cytosol causing calcium overload during pacing.

**Fig. 6.**
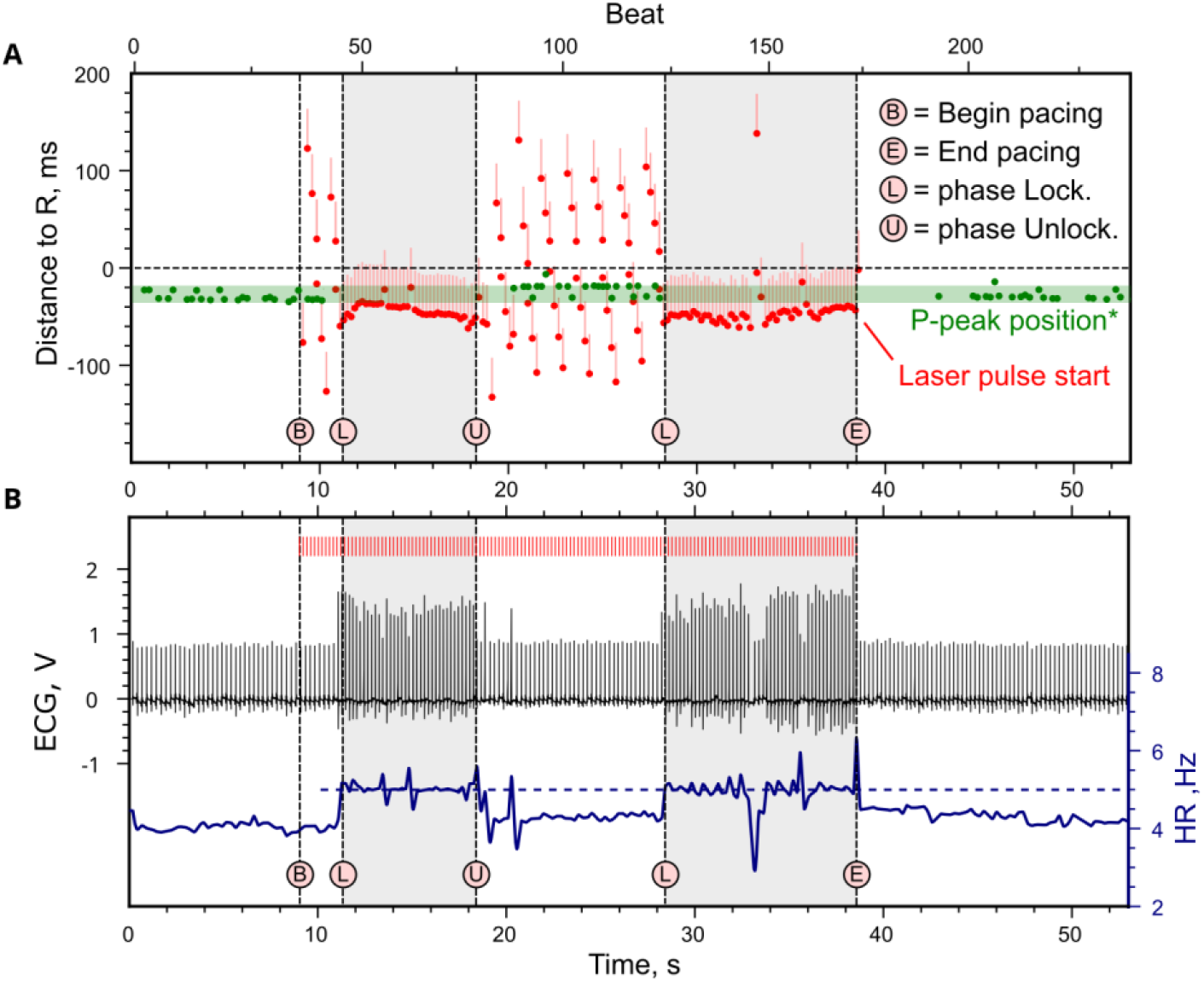
Loss and recovery of phase locking during left ventriculum pacing with 5-Hz 20-ms trigger pulses. **a**, Position of the P-peak and laser pulse in relation to the R-peak in ms. **b**, Change of complex shape (black curve) and heart rate (blue curve) during the experiment. Vertical red lines demonstrate trigger pulses. Critical events are marked with ‘B’/’L’/’U’/’E’ letters (see the legend). **\*** In this sample the atrium beat was not synchronized with laser pulses that is why it was hard to separate P-peaks from the noise during the period of synchronization.

**Fig. 7.**
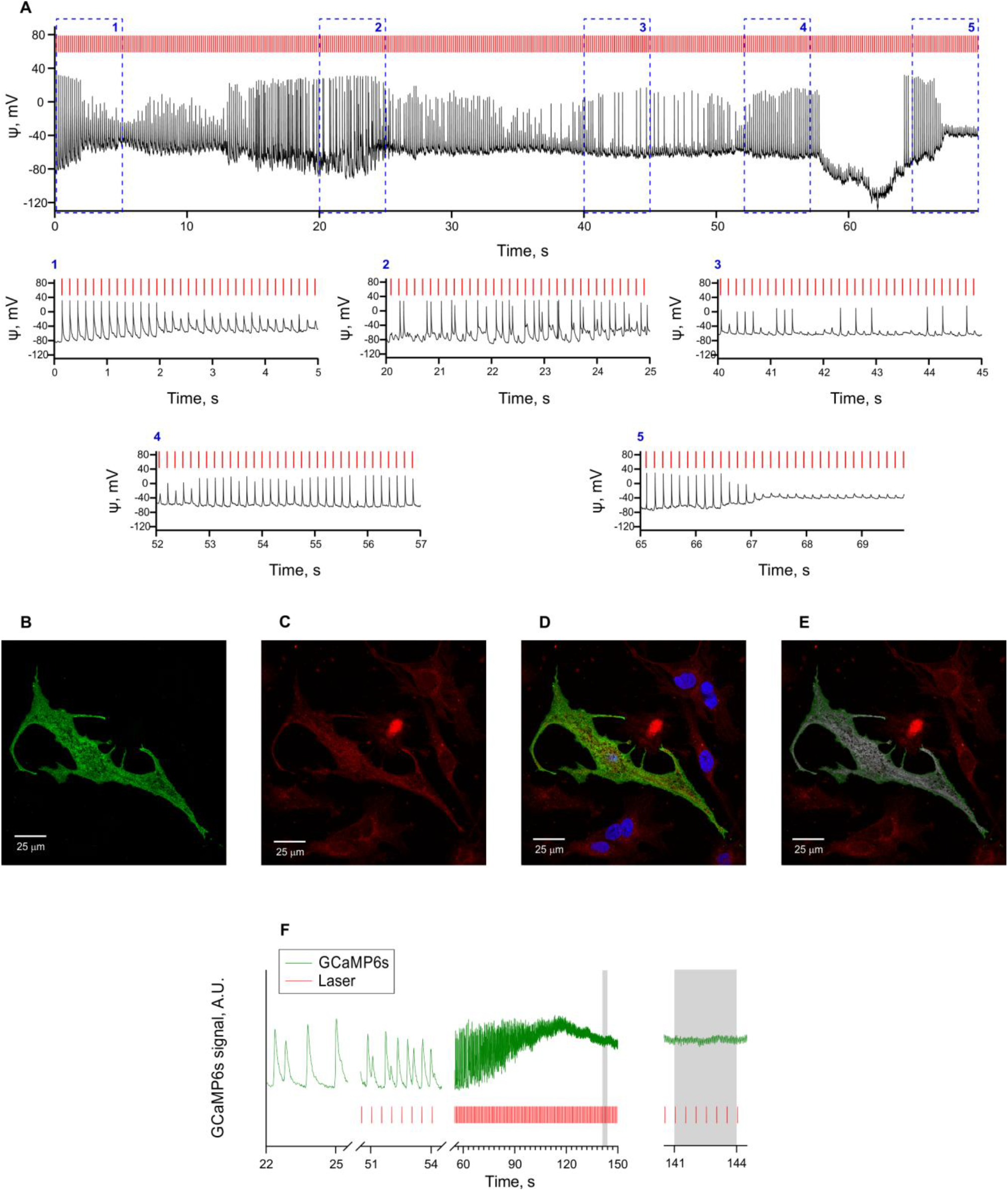
Long pacing, hTRPV1(−109aa) localisation and calcium dynamics upon heating in murine neonatal cardiomyocytes. **a**, Action potentials of an isolated neonatal murine cardiomyocyte during long pacing. **b–e**, Cellular localization of hTRPV1 in transduced neonatal murine cardiomyocytes. **b**, Staining of hTRPV1-FLAG with anti-FLAG antibodies. **c**, Staining of ER with ER-tracker Red. **d**, Merged image. **e**, Colocalization of anti-FLAG and ER-tracker staining (shown in greys). **f**, Calcium pulses in a neonatal murine cardiomyocyte transduced with hTRPV1(−109 a.a.) and heated by IR laser with a repetition rate of 2 Hz and 30 ms pulse width in calcium-free medium.

To check if the hTRPV channels in ER are functional and if their activation can induce an increase in cytosolic Ca^2+^ level, we placed neonatal murine cardiomyocytes transduced with TRPV1(−109 a.a.) in a calcium-free medium and applied the specific TRPV1 activator capsaicin. TRPV1(−109 a.a.)-expressing cells demonstrated a significant calcium response to capsaicin **(Supplementary Fig. 15)**. Since calcium is absent in the extracellular medium, the ER is the most probable source of calcium under these conditions.

### Method safety

The possible calcium overload during thermogenetic pacing raises questions about the pro-arrhythmogenic effect of this kind of stimulation. To check this, we induced calcium overload in isolated hearts by perfusing them with Tyrode’s solution supplemented with 1 mM of caffeine. Under these conditions, 1:1 phase lock, once established, was lost in a few seconds, turning into 2:1. However, we did not observe re-entry waves or other events that could be considered pro-arrhythmogenic **(Supplementary Fig. 16, Supplementary Video 1)**.

Another question comes from the data demonstrating small but detectable inward currents through TRPV1 channels at temperatures between 38°C and 41°C which can occur in humans e.g. during infectious diseases. We investigated the effect of TRPV1 hyperexpression under hyperthermic conditions. Mice were anesthetized with isoflurane, breathed spontaneously, and were thermally insulated. After that, the pad temperature was slowly raised to achieve a rectal temperature above 43°C. All the mice died from respiratory arrest but did not demonstrate arrhythmias. We did not find significant differences in heart rate variability and the shape of QRS complexes between mice expressing TRPV1 and control animals **(Supplementary Table 3)**. We hypothesize that cardiomyocytes have enough ion pumps to resist raising TRPV1-dependent calcium influx if the heating is slow.

Since the expression of TRPV1 channels has been demonstrated in other cell types located in the heart (e.g., fibroblasts, endothelial cells, smooth muscle cells, and autonomic neurons), continuous heating may theoretically be harmful to these cells. We tried to investigate possible off-target effects using non-myocardial cells isolated from neonatal murine hearts. We plated these cells, loaded them with a fluorescent Ca^2+^ dye Fura-4 AM, and heated them with a nichrome wire loop. Non-myocardial cells did not demonstrate any significant calcium dynamics **(Supplementary Fig. 17)** suggesting that TRPV1 channels in these cells, if present, do not generate big inward currents.

## Discussion

In the present study, we demonstrated that thermogenetics can be applied for cardiac pacing. This technology opens the way for the search of new heart rhythm control and defibrillation strategies.

Thermogenetics is quite similar to optogenetics but utilizes ion channels activated by temperature instead of light. In previous studies, optogenetics was successfully used to control the heart muscle *in vitro* and *in vivo*. In 2010, the possibility of heart rhythm control in mammals using transgenic mice expressing channelrhodopsin-2 was demonstrated^10^. Five years later, two groups demonstrated the possibility of using AAVs for the channelrhodopsin-2 gene delivery to the heart^11,12^. Over the next ten years, cardiac optogenetics developed extensively, which led to the creation of “all-optic cardiac electrophysiology”^30^. Nevertheless, the major problems of optogenetics — the low penetrance of the visible light and the immunogenity of opsins — remained unsolved.

Thermogenetics uses temperature sensitive channels which are expressed in non-immune-privileged tissues. We have shown that human TRPV1 can be used as a molecular tool for this technology, which opens possibilities for potential therapeutic methods in various fields of medicine in the future. TRPV1 activation was confirmed using the patch clamp method, as well as the genetically encoded fluorescent indicator GCAMP6S, which visualizes an increase in calcium flux. This channel is well-characterized and was successfully used in several studies for brain stimulation with magnetic fields heating nanoparticles in tissue^27,31,32^.

TRP channels can be activated in various ways: by using chemical agonists such as capsaicin, by changing the ambient temperature^33-35^ causing a change in the temperature of the whole animal, or by using IR laser stimulation^22,23,36^. In our study, the use of an IR laser setup made it possible to achieve a high spatial resolution and a fast channel activation rate. However, IR lasers have significant limitations. First, the non-invasive pacing cannot be achieved, because the radiation should be directly delivered to the tissue by an optical fiber. Second, IR light although penetrates deeper than e.g. blue light heats only the superficial tissue layer. We believe that high-intensity focused ultrasound (HIFU) is a much more promising technique for non-invasive heating of tissues. Depending on the transducer size and focus, HIFU allows to heating of big volumes deep in tissues. We performed some tests with a setup including a HIFU transducer coupled with an ultrasound probe for navigation and successfully heated tissues in 6×1 mm (depth × diameter, approximately 3.1 mm^3^) ellipsoid volume. However, our current setup is too large and does not provide enough resolution to navigate to a mouse heart, so we did not use it in the present study. Recent studies demonstrate the possibility of heating freely moving mice’s brains by HIFU^37,38^. This technique may make truly non-invasive heart pacing possible, but it must be considered that in addition to heating, this technology includes cavitation and mechanical effects at certain powers and frequencies^39^.

Excessive intake of calcium into the cell cytosol from the ER during stimulation of cardiomyocytes with the IR laser is an important factor limiting the duration of thermogenetic pacing. As a result, the pacing is short-term, however, we suppose further studies can help find ways to increase the pacing duration. A possible way to overcome this limitation is to use engineered TRPV1 channels with lower calcium conductance and lower ER tropism. Such channels will cause cell depolarization primarily due to the sodium inward current, preventing calcium overload. Another way could be to use multipoint laser heating with several pacing initiation sites in different areas of the heart. In this case, the points would be heated sequentially over a short time interval. This approach will avoid calcium overload of each individual point due to the additional rest time obtained and the normalization of calcium concentration in the cytosol. Another possible way to overcome current limitations is to target TRPV1 to specific cell types, such as pacemaker cells^40^, Purkinje fibers^41^, or cardiac ganglia^42^. The use of one of the proposed approaches or their use in combination may contribute to longer and more effective pacing.

The current setup cannot guarantee stable and complete 1:1 phase locking during pacing probably because of the effects described above. It occurs when the stimulation strength at the available maximum appears to be still weak. Interestingly, this “weak” stimulation sometimes bears resonance phase locking of 4:3 / 3:2 which was not previously described in the literature. Weak electric stimulation also produces resonance locks^43,44^ but their numbers are different because the mechanism of stimulation is different. This phenomenon itself is very interesting to study in the context of arrhythmia genesis.

We performed studies on the safety of the method and did not find any pro-arrhythmogenic effects of the thermogenetic pacing. However, mice are not an optimal model for studying arrhythmias. The electrophysiology of the heart in mice differs from larger mammals, including humans, which is another limitation of the present work. Nevertheless, the mice’s heart rate is higher, so in humans, heart tissue has more time for cooling down and Ca^2+^ efflux from the cytosol. Since methods for gene delivery into myocardium are developing rapidly, the thermogenetic platform we present is capable of scaling to larger animals. This allows further research which is needed to establish the clinical perspectives of the method. Another safety consideration considers the possible off-target effects on the cells endogenously expressing TRPV1 channels. Although we did not demonstrate any reaction of cardiac cells isolated from murine neonatal hearts, long-term studies are needed to confirm the absence of any side effects. TRPV1 should be much less or rather even completely not immunogenic because it is a human protein naturally expressed in many tissues, unlike, for example, the light-sensitive channels used for optogenetics. Genetic modification of cardiomyocytes has its own risks, but the safety of these methods is continuously being improved, and it is probably not necessary to modify the whole heart. This question will be further addressed during preclinical studies. It was demonstrated that AAVs are suitable for long-term genetic modifications of tissues, including myocardium: after a single intracoronary infusion of AAVs clinical effects persist for at least 3 years^45^, and e.g. in skeletal muscles the transgene expression may be stable for 10 years^46^.

In the present study, we used thermogenetics for heart pacing. It was demonstrated before that optogenetics can be used for the termination of ventricular arrhythmias in the whole heart^47^. Similar use of thermogenetics can help to develop methods for non-invasive and painless cardioversion.

This is the first study on cardiac thermogenetics. We believe that our approach holds promise not only for basic research but also for translational medicine.

## Methods

### Genetic constructs and viruses

In the present work, we used adeno-associated viruses (AAVs) serotype PHP.S^48^ and DJ serotype^49^ for cTnT_gCaMP6s construction. The construction pUCmini-iCAP-PHP.S for AAV-PHP.S serotype assembly was kindly gifted by Viviana Gradinaru (Addgene plasmid #103006). For targeted expression of constructs in cardiomyocytes, we used a specific promoter — a constitutive hybrid promoter composed of the CMV immediate-early enhancer fused to the cardiac troponin T (cTnT) promoter. The plasmid containing cTnT was kindly gifted by Thomas Michel; Addgene plasmid #119164. pAAV cTnT_hTRPV1 (−109 amino acids residues) P2A mRuby2 (P2A — a self-cleaving peptide), pAAV cTn_ hTRPV1-3×FLAG tag and pAAV cTnT_GCaMP6s were constructed for this work. All genetic constructs were assembled by AQUA-cloning ^50^. pAAV cTnT HyPerDAAO plasmid was cleaved with SacI (Thermo Scientific, FD1133) and BglII restriction enzymes (Thermo Scientific, FD0083) to remove HyPerDAAO. For further assembly, the linearised vector was used. Genes encoding GCaMP6s and mRuby2 were amplified from plasmids previously generated in our laboratory.

The creation of short hTRPV1 (−109 amino acid residues) was based on the available pCAG_hTRPV1-p2A-tdTomato construct. The DNA sequence between the XbaI (Thermo Scientific, FD0684) and XhoI (Thermo Scientific, FD0695) restriction sites, with fragments of the chimeric intron and channel sequence, was inserted into the pAL2-T vector (Evrogen, TA002). Using the Tersus Plus PCR kit (Evrogen, PK221), blunting of the protruding ends in combination with A-tail treatment was performed. The overall construct was used to generate several short channel variants, including pCAG_(−109aa)hTRPV1-p2A-tdTomato. Then, a PCR reaction was performed. The PCR product was extracted using the Cleanup Standard kit (Evrogen, BC022S), and then treated with T4 polynucleotide kinase (Thermo Scientific, EK0031) and ligated. As a result, the sequence containing the chimeric intron and truncated channel fragments was cloned from the initial pAL2-T vector into pCAG_hTRPV1-p2A-tdTomato using XbaI and SmaI (Thermo Scientific, FD0663) restriction sites.

The genetic constructs pCAG_hTRPV1_p2A_GCaMP6s and pCAG_hTRPV1-3×FLAG_p2A_GCaMP6s for transfection of the HEK 293 cell line were assembled using the AAV-CAGGS-insertion plasmid available in the laboratory. The plasmid was linearized by MunI (Thermo Scientific, FD0754) and MluI (Thermo Scientific, FD0564) restriction enzymes. Genes encoding hTRPV1, hTRPV1-3×FLAG, GCaMP6s, and a self-cleaving peptide P2A were amplified from available plasmids using Q5 High-Fidelity DNA Polymerase (M0491L) and extracted using the Cleanup Standard kit (Evrogen, BC022S). The DNA fragments were inserted in the linearized AAV-CAGGS-insertion plasmid with ClonExpress MultiS One Step Cloning Kit (Vazyme, C113-01).

Construct verification was performed by sequencing. Midiprep was prepared using the Plasmid Midiprep 2.0 kit (Evrogen, #BC124) according to the manufacturer’s instructions.

The 5′-end of primers was homologous to the vector and the 3′-end was specific to the particular sequence. The homemade chemically competent *Escherichia coli* Top10^51^ strain was used for the cloning, maintenance, and propagation of plasmids. Overall, the accurate assembly of the constructs and absence of mutations were verified by sequencing (Evrogen). Midipreps were prepared using the Plasmid Midiprep 2.0 kit (Evrogen, #BC124) according to the manufacturer’s instructions. Then, the constructs for gene delivery to in vivo and in vitro cardiomyocytes were packaged into AAV viral particles at the Viral Core Facility of Shemyakin-Ovchinnikov Institute of Bioorganic Chemistry. Virus titers were 2.1·10^12^ and 6.2·10^12^ VG/ml for pAAV_cTnT_hTRPV1(–109 amino acids residues)_P2A_mRuby2, 2.1·10^12^, 1.6·10^12^, 8.1·10^12^, 6.1·10^12^, 9.1·10^12^ VG/ml for pAAV_cTnT_hTRPV1-3×FLAG, 3.6·10^12^ for pAAV_cTnT_gCaMP6s.

### Cardiomyocyte cell cultures and transduction

The mixed mouse primary neonatal cardiomyocyte cell culture was obtained using a neonatal heart dissociation kit (Miltenyi Biotec, 130-098-373) according to the manufacturer’s instructions. The cells were cultured in Dulbecco’s Modified Eagle’s Medium/Nutrient Mixture F-12 Ham (DMEM/F12), 1:1 mixture (BioloT, 1.3.7.2.) supplemented with 10% fetal bovine serum (FBS, Biosera, FB-1001/500), penicillin 100 U/ml /streptomycin 100 mg/ml (PanEko, A065Π), and L-glutamine 0.365 g/l (PanEko, Ф032). For electrophysiological experiments, 5.5*104 cells were seeded onto 10mm microscope glass coverslips (Heinz Herenz, 1051199), which were coated with 10 mg/ml gelatin from bovine skin (Sigma-Aldrich, G9391-100G) diluted in phosphate-buffered saline (PBS, PanEko, P071-1/B-60201) and maintained at 37°C in 5% CO^2^. For transient expression of the hTRPV1 channel, a reporter protein, and a fluorescent Ca^2+^ sensor GCaMP6s, we used AAV-based vectors with the encoded genes above. For infection of the cardiomyocytes, we used the AAV-DJ serotype at a MOI of 12,000 VG/cells for cTnT_hTRPV1(sh)_P2A_mRuby based viruses, and a MOI of 2,500 VG/cells for cTnT_GCaMP6s ones. The cells were infected on the next day after plating, and the transgene expression peak was observed on the third day after the infection.

As optical mapping requires a dense layer of functional cells, a suspension of isolated cells was seeded at a concentration of 2.5*10^5^ cells/cm^2^ on 25mm microscope glass coverslips (Heinz Herenz, 1051199). The coverslips were coated with 0.16 mg/mL fibronectin (Imtek, H Fne-C) diluted in phosphate-buffered saline (PBS, PanEko, P071-1/B-60201) and maintained at 37°C in 5% CO^2^ for 12 hours before seeding. For transient expression of the hTRPV1 channel, we used AAV-DJ virus with cTnT_hTRPV1-3×FLAG construct at a MOI of 100,000 VG/cells. The cells were infected on the next day after plating, and the transgene expression peak was observed on the third day after the infection.

### HEK 293 cell culture and transfection

HEK 293 were cultured in Dulbecco’s Modified Eagle’s Medium/Nutrient Mixture F-12 Ham (DMEM/F12), 1:1 mixture (BioloT, 1.3.7.2.) supplemented with 10% fetal bovine serum (FBS, Biosera, FB-1001/500), penicillin 100 U/ml /streptomycin 100 mg/ml (PanEko, A065Π), and L-glutamine 0.365 g/l (PanEko, Ф032). Cells were seeded onto 10mm microscope glass coverslips (Heinz Herenz, 1051199), which were coated with 10 mg/ml gelatin from bovine skin (Sigma-Aldrich, G9391-100G) diluted in phosphate-buffered saline (PBS, PanEko, P071-1/B-60201) and maintained at 37°C in 5% CO2. At least 18 h prior to the experiment, cells at approximately 60% confluence were transfected with plasmid construct pCAG_hTRPV1_p2A_GCaMP6s or pCAG_hTRPV1-3×FLAG_p2A_GCaMP6s using GenJect reagent (Molecta, Gen39-1000p) according to the manufacturer’s protocol.

### solation of non-cardiac cells

The non-myocyte cell fraction was obtained by dissociation of neonatal mouse hearts using a neonatal heart dissociation kit (Miltenyi Biotec, 130-098-373) according to the manufacturer’s instructions. The mixed cell suspension obtained from the dissociation was separated into cardiomyocytic and non-cardiomyocytic fractions according to the following protocol. The cell suspension of 8 hearts was resuspended in 2 ml 1 x ADS buffer (116 mM NaCl, 20 mM HEPES, 1 mM NaH2PO4, 5.6 mM D-glucose, 5.4 mM KCl, 0.8 mM MgSO4, pH = 7.35) and separated in a Percoll (Cytiva, 17089101) density gradient. Stock Percoll was composed of 90% percoll and 10% 10 x ADS buffer. To prepare the gradient, 1 part of the Stock Percoll was diluted with 1.2 parts of 1 x ADS to prepare top Percoll with a low density (1.060 g/ml) and 0.5 parts of 1 x ADS to prepare bottom Percoll with a high density (1.086 g/ml).

Subsequently, 3 ml of bottom percoll, 4 ml of top percoll, and 2 ml of cell suspension were added to a 15-milliliter conical tube and centrifuged at 1500g for 30 min with no deceleration brake at 4 °C. Non-cardiac cells residing at the border of ADS and top percoll were selected into a 15-ml tube and washed twice with DMEM/F12 medium (BioloT, 1.3.7.2.).

The collected cells were seeded onto confocal dishes (SPL Life Sciences, 101350) which were coated with 10 mg/ml gelatin from bovine skin (Sigma-Aldrich, G9391-100G) diluted in phosphate-buffered saline (PBS, PanEko, P071-1/B-60201) and maintained at 37°C in 5% CO^2^. The cells were cultured in Dulbecco’s Modified Eagle’s Medium/Nutrient Mixture F-12 Ham (DMEM/F12), 1:1 mixture (BioloT, 1.3.7.2.) supplemented with 10% fetal bovine serum (FBS, Biosera, FB-1001/500), penicillin 100 U/ml /streptomycin 100 mg/ml (PanEko, A065Π), and L-glutamine 0.365 g/l (PanEko, Ф032).

### TRPV1 channel localization in neonatal mice cardiomyocytes

To understand the localization of the expressed TRPV1 channels in neonatal mice cardiomyocytes, we utilized cells infected with AAV-PHP.S serotype viruses with pAAV_cTnT_hTRPV1_3×FLAG at a MOI of 2500 VG/cells. To visualize the endoplasmic reticulum, we stained the live cells with ER-Tracker™ Red (BODIPY™ TR Glibenclamide, Thermo Fisher Scientific, E34250) according to the manufacturer’s instructions. After the staining with ER-tracker, cardiomyocytes were fixed with 4% paraformaldehyde (Sigma-Aldrich, 158127-100G) for 5 minutes at room temperature and washed trice with 0.3% Tween 20 (Sigma-Aldrich, P1379-250ML) diluted in PBS (5 minutes each). The fixed cells then were blocked with PBS containing 0.12% tween 20, 1% bovine serum albumin (BSA, PanEko, 68100.10Г), and 10% goat serum (Thermo Fisher Scientific, 16210072) for 40 minutes at room temperature. After the buffer removal, the cardiomyocytes were labelled with DYKDDDDK Tag Recombinant Rabbit Monoclonal Antibody (8H8L17, Invitrogen, MA1-142-A488) at 1:500 dilution in 1% BSA, 10% goat serum, and 89% PBS for 2 hours at room temperature. The samples were washed three times with PBS after the incubations. Then, the cells were stained with Goat anti-Rabbit IgG (H+L) Cross-Adsorbed Secondary Antibody Alexa Fluor™ 488 (Invitrogen, A-11008) at dilution 1:500 for 1 hour at room temperature. The removal of non-conjugated antibodies was performed in parallel with cell nuclei staining.

The cells were incubated with PBS supplemented with 2 μg/ml DAPI (Miltenyi Biotec, 130-111-570) for 15 minutes at room temperature. For further experiments, glasses with the labelled cells were placed onto the Superfrost Plus adhesion slides (Epredia, EPBRSF41296SP) in 20 µl VECTASHIELD Vibrance Antifade Mounting Media (Vector Laboratories, H-1700-2) and stored at +4°C in the dark.

The samples were analyzed using an inverted Nikon A1 confocal microscope and visualized using Nikon NIS-Elements software. We pictured the sample in each channel individually exciting DAPI, DYKDDDDK Tag-Alexa Fluor 488, and ER-tracker by laser lines 405 nm, 488 nm and 561 nm, respectively. Colocalization analysis was performed using ImageJ software.

### Optical mapping in cell culture

Optical mapping with the Ca^2+^ chemical indicator Fluo4-AM (Lumiprobe, 1892-500ug) at a concentration of 4 ug/ml was performed in Tyrode’s solution (pH 7.25-7.4) according to the protocol described previously in^52^. The signal was recorded at a resolution of 64×64 pixels and a sampling rate of 239 frames per second (Olympus MVX-10 Macro-View fluorescence microscope (Olympus Co., Tokyo, Japan), Andor iXon-3 EMCCD Camera (Andor Technology Ltd., Belfast, UK) high-speed camera). Data were recorded at 37°C using both thermal and electrode stimulation. The duration and amplitude of the electrode and thermal stimulation ranged from 1 ms to 20 ms duration and from 1 V to 6 V. The stimulation frequency ranged from 2 to 4 Hz unless otherwise stated according to the stimulation protocols. The stimulus was set using a generator (2 MHz USB PC Function Generator, PCGU100 (Velleman, Gavere, Belgium)). Platinum electrodes were used.

Data processing was performed using ImageJ software. The ImageJ plug-in (time-lapse color-coder) was used to construct pseudo-three-dimensional images and activation maps. Principal component analysis was performed in Wolfram Mathematica 12.

### Electrophysiology of single cardiac cells

Patch electrodes were pulled from hard borosilicate capillary glass (Sutter Instruments flaming/brown micropipette puller) and filled with an intracellular solution consisting of (in mM) K-gluconate, 100; KCl, 40; HEPES, 10; NaCl, 8; MgATP, 4; MgGTP, 0.3; phosphocreatine, 10 (pH 7.3 with KOH) in some experiments EGTA (15mM) was added to the intracellular solution. Cells were identified visually using IR-video microscopy using a Hamamatsu ORCA-Flash4.0 V3 Digital sCMOS camera (Hamamatsu Photonics) expression of wild type or mutant TRPV1 in the cardiomyocytes was confirmed by the presence of RFP. Coverslips were placed in a recording chamber continuously perfused with heated Tyrode’s solution. Whole-cell recordings were taken at 32°C in current-clamp mode using a HEKA EPC-10 amplifier (List Elektronik) with a sampling rate of 100 μs. Steady state current was injected to achieve a membrane potential of approximately -60 to −90 mV. For experiments with capsaicin, it was applied directly to the cell via micropipette. For experiments using the IR laser, the optic fibre was placed near the cell, and light from a green laser diode was shone onto the cell to check correct positioning. The laser was controlled via the TTL output from the HEKA EPC-10 amplifier. Laser intensity was chosen to achieve the desired temperature.

The parameters of the generated action potentials were calculated from the raw records. The amplitude was calculated as the level of the depolarization local peak with respect to the base level. The rate of depolarization and repolarization were estimated as the first-time derivative of the potential. Action potential duration was calculated as the width of the signal at the level of 50% of the amplitude (half-width). A two-way rank sum test was used in the statistical evaluation of the samples. Differences with a probability level p<0.05 were considered as significant.

### Electrophysiology of HEK 293 cells

The calibration of temperatures attained at specific IR laser diode currents was conducted by monitoring the currents flowing through an open microelectrode. Tyrode’s perfusion solution was heated to 50°C as measured by a thermistor in the recording chamber, and then the solution was cooled while recording the current through the open microelectrode. The temperature was plotted against the current through the open electrode to create a calibration curve. Subsequently, the same open microelectrode was heated via IR laser at different intensities, and stable currents were measured. Utilizing the established calibration curve and the currents measured during the heating process by the IR laser, an approximation of the temperature for varying laser intensities was obtained.

HEK cells were patched on the same setup and with the same intracellular solution as described in «Electrophysiology of single cardiac cells» at temperature 32-33°C. Cells expressing TRPV1 or TRPV1_3×FLAG were identified by the presence of GCaMP6s using a Hamamatsu ORCA-Flash4.0 V3 Digital sCMOS camera (Hamamatsu Photonics). The cells were held in a voltage clamp at -20mV and heated by an IR laser, with each step increasing the IR intensity. The resulting currents were recorded.

### Intracellular calcium recordings of single cardiac cells

Neonatal cardiomyocyte cells transduced with GCaMP6s sensor were viewed and acquired under a water immersion Olympus LUMPLFLN40×W objective with 40X magnification. Data acquisition was performed at 20 fps using a Scientifica SliceScopePro 2000 microscope (Scientifica, UK) equipped with a Hamamatsu Orca Flash 4.0 CMOS monochrome digital camera (Hamamatsu Photonics) connected to a PC running the free software uManager. A CoolLED pE-300ultra was used as a light source. It was synchronized with the laser heating system via BNC-TTL output from the Heka Elektronik EPC 10 USB Patch Clamp Amplifier. The GCamp6s signal was analyzed using Fiji software.

### Intracellular calcium recordings of non-cardiomyocyte cells

In order to record intracellular calcium dynamics in nonmyocytes cells upon heating, the growth medium was first withdrawn, and cells were placed in HBSS solution (PanEko, CP020-50) with Fluo4-AM calcium indicator (Lumiprobe, 1892-500ug) at a concentration of 2.8 ug/ml for 30 minutes. The cells were then placed in an incubator at 37°C. Subsequently, the dye solution was replaced with pure HBSS buffer. Cells were heated according to the previously described protocol^53^. Data processing was performed using ImageJ software.

### Mice

Experiments were carried out using C57Bl/6J mice (The Jackson Laboratory, #000664, RRID: IMSR_JAX:000664) in compliance with all the ethical regulations for work with experimental animals and according to the European Convention for the Protection of Vertebrate Animals used for Experimental and other Scientific Purposes (1986, ETS 123). The animals were bred and housed in the animal facilities of the Institute of Bioorganic Chemistry of the Russian Academy of Sciences (IBCh RAS) in a 12 h light-dark cycle with free access to food and water. All procedures were approved by IBCh IACUC protocols No. 356 and No. 358.

### Electrocardiography of immobilized mice

The principal scheme is illustrated in **Supplementary Fig. 18**. The setup consisted of a laser source (assembled as described in section “Distant heating systems”), a JDS6600 electrical stimulator (Simac Electronics, Germany), external 10-MHz ADC E20-10 (L-CARD, Moscow, Russia), an IT-24P ultra-fast type T thermocouple (Physitemp, NJ, USA), an external High-Pass filter (handmade), two DPA-2FS biopotential amplifiers (NPI Electronics, Germany), one LP-04-M fixed-gain preamplifier (L-CARD, Moscow, Russia), a PC with PowerGraph 3.x preinstalled (DiSoft, Moscow, Russia), a homeothermic monitoring system (ThermoStar, China), and a SAR-1000 small animal ventilator(CWE, USA).

The ECG was recorded in a 4-electrode scheme (‘R’, ‘L’, ‘RL’, and ‘LF’). The anaesthetized mouse was immobilized onto a homeothermic monitoring system, and F9049 disposable sticky electrodes (FIAB, Italy) were mounted to fix each paw on the mat. Each paw was preliminarily treated with keratinase (VEET, France) for better electrical contact. For recording, we used two differential amplifiers: we fed the R and L leads to their non-inverting inputs, and the LF lead to the inverting inputs. The RF lead was connected to the ground of the amplifiers, as well as to the screen of the foil surrounding the mouse from the bottom and sides. High-pass and low-pass frequencies were set to 0.3 and 10 kHz respectively, 100x gain was used. The output signal was additionally processed with a 0.1 Hz High-pass filter. The E20–10 ADC was operated with a 25 kHz sampling frequency mode per channel and 12-bit resolution. PowerGraph was set to remove 50-Hz interference from the mains using a digital band-cut filter (48–52 Hz) from both ECG signals (R–LF and L–LF). After band cutting, PowerGraph was instructed to subtract one input from another to calculate the R–L ECG signal, which was visualized and further saved to CSV files for programmatic analysis (see “Raw ECG data analysis”).

To synchronize laser pulses with the ECG signal, we operated the laser diode driver (LaserSource 4320) through its modulation input with the electrostimulator and fed the same trigger pulses to the separate channel of the ADC (**Supplementary Fig. 18, IN_4**). To prevent the stimulator from interfering with the recorded signal, it was not connected to the main grid and received power from a powerbank staying galvanically isolated. The LD driver provides a time delay of less than 1 µs between the trigger pulse and operational current pulse, which is much lower than the time scale of the observed effects.

To control the temperature with high time resolution, we used an ultrasmall IT-24P T-couple (time constant 4 ms). This sensor was connected to a differential preamplifier (**Supplementary Fig. 18**, LP-04-M). We did not use cold junction compensation for external temperature correction; therefore, we calibrated our thermocouple before every measurement, using a solid-state thermostat. To avoid possible negative effects caused by the thermocouple implantation, we only recorded temperature in control mice. We used the same IR laser modes for experimental and control animals. Body temperature was maintained at 38°C with a thermal pad coupled with a rectal thermometer.

### Viral vector delivery

The viral vectors were delivered to mice hearts by an injection into the jugular vein. 6– 8-week-old female mice were anaesthetized with 5% isoflurane (Baxter, 10019036040) in an induction chamber (SomnoSuite, Kent Scientific), and then placed in the supine position onto a heating pad (Digital stereotaxic instrument, Stoelting Co). For deep anesthesia, 1.5% isoflurane was administered through an inhalation mask during the operation. The hair was removed from the thorax with a trimmer, and the operating field was treated with 70% ethanol. Prior to incision, 0.25% of local anaesthetic bupivacaine (Ozon) was injected into the area lateral to the midline of the body. After several minutes, a small incision was made from the pectoral muscle to the lower part of the neck. Adeno-associated viruses were injected into the jugular vein using insulin syringes with a 27-gauge needle. Each animal was injected with approximately 100 μl of a suspension of pAAV_cTnT_hTRPV1_3×FLAG AAV-PHP.S serotype viral particles for a total of 6-9·10^11^ genomes per mouse. The wound was sutured, and the mice were intraperitoneally injected with ketoprofen (5 mg/kg, Sandoz).

### Immunohistochemistry

Expression of TRPV1 channel in tissues after intravenous injections of pAAV_cTnT_hTRPV1_3×FLAG in mice was confirmed by immunohistochemical staining of slices. The mice were deeply anaesthetized with 3% isoflurane and decapitated, the hearts were isolated and placed in Tissue-Tek Cryomold (Sakura Finetek, 62534-10) filled with Tissue-Tek O.C.T. Compound (Sakura Finetek, 4583). The mould with the heart was frozen in liquid nitrogen vapor and then stored at −80°C or immediately dissected into thin 15-30 μm sections using an HM525 NX cryostat (Thermo Scientific) at −16°C. The prepared samples were placed onto the Superfrost Plus adhesion slides and stored at −20°C.

Heart tissues were fixed in 4% paraformaldehyde in PBS for 5 minutes at room temperature and then washed with 0.3% Tween 20 in PBS three times with further incubation for 5 minutes. To block nonspecific binding, slices were incubated in a blocking buffer (10% goat serum, 1% bovine serum albumin, 0.15% Tween 20, PBS) for 1 hour at room temperature, then washed again with 0.3% Tween 20, PBS. Staining with primary antibody (DYKDDDDK Tag Recombinant Rabbit Monoclonal Antibody (1:500, 8H8L17, Invitrogen, 701629), Anti-HCN4 Antibody (1:200, Polyclonal, Alomone Labs, APC-052-GP)) was performed in blocking solution at 4°C for 14 hours or overnight. Slices were then washed three times with 0.3% Tween 20, PBS. Incubation with secondary antibody (Goat anti-Rabbit IgG (H+L) Alexa Fluor™ 647 (1:500, Invitrogen, A-21244) Goat anti-Rabbit IgG (H+L) Alexa Fluor™ 488 (1:500, Invitrogen, A-11034), Goat anti-Guinea Pig IgG (H+L) Alexa Fluor™ 647 (1:500, Invitrogen, A-A-21450)) was also performed in a blocking buffer for 2 hours at room temperature. The stained samples were incubated for 15 minutes in PBS supplemented with 2 μg/ml DAPI and washed by distillate water once. The slides with the stained slices were covered with 10mm microscope glass coverslips with 20 µl VECTASHIELD Vibrance Antifade Mounting Media. The specimens were stored in the dark at +4°C.

The localization of hTRPV1 and HCN4 was analyzed by confocal microscopy via an inverted Nikon A1 confocal microscope and visualized using Nikon NIS-Elements software. For DAPI, Alexa488, and Alexa647 excitation, 405 nm, 488 nm, and 633 nm lasers were used accordingly.

### TRPV1 channel distribution in the mouse heart

To determine the number of transduced cardiomyocytes sufficient to induce cardiac pacing, we conducted a quantitative analysis of FLAG immunopositive cells in hearts expressing the hTRPV1-3×FLAG protein. Thus, we obtained micrographs from 10 slices per mouse: 3 from the atria, 4 from the ventricles, and 3 from the apical part of the cardiac myocardium. Each image was 420 x 420 μm in size. The resulting images were analyzed using the image processing software Fiji. Cells immunopositive for FLAG were manually selected and taken for statistics. Cells without staining were also selected to obtain the percentage of transduced cells to non-transduced cells in this area.

To visualize the intracellular distribution of TRPV1 channels, we obtained Z-stacks of immunopositive FLAG+ cells from heart slices expressing TRPV1-3×FLAG. Images were acquired using an inverted Nikon A1 confocal microscope equipped with a Nikon CFI Plan Lambda D 60x/1.42na Oil Objective. Each Z-slice was 0.5 μm in size. The subsequent processing of the data was facilitated by the utilization of Nikon NIS-Elements software.

### Mice heart perfusion and optical mapping

The mouse was anesthetized using an anesthesia machine (R550 Multi-output Animal Anesthesia Machine (RWD, Shenzhen, China)) and decapitated. The heart was isolated according to the previously described protocols^54,55^ and then rinsed with 37°C Tyrode’s solution (Sigma-Aldrich, T2145) and heparin (Belmedpreparaty, B01AB01) at a concentration of 50 ME/ml. Staining with the Ca^2+^ chemical indicator Fluo4-AM (Lumiprobe, 1892-500ug) at a concentration of 2.8 ug/ml as well as optical mapping were performed in Tyrode’s solution at 37°C. Heart fixation to the cannula through the aorta was performed using surgical filament. The time elapsed from the moment of mouse decapitation to the start of perfusion through the cannula was less than 10 minutes. During staining and optical mapping, the heart was perfused using a Langendorff apparatus. The setup consisted of a perfusion circuit and a recording optical system based on a high-speed optical mapping unit (Olympus MVX-10 Macro-View fluorescence microscope (Olympus Co., Tokyo, Japan), Andor iXon-3 Camera 897-U high-speed camera (Andor Technology Ltd., Belfast, UK)). The perfusion circuit was designed to maintain a fixed fluid volume. It consisted of a peristaltic pump (Masterflex L/S Digital Drive, 600 rpm; 115/230 VAC, Masterflex L/S Easy-Load® II Pump Head, SS Rotor; 2-Channel (Cole-Parmer Instrument Company)), a thermostat (Cole-Parmer Polystat Standard 6.5 L Heated Bath, 150 C, 115 VAC/60 Hz, (Cole-Parmer Instrument Company)), oxygenator (Cole-Parmer Bubble Latcher, Water Shirted Reservoir, Oxygenating Bubbler (Cole-Parmer Instrument Company)). The total volume of fluid circulating in the device was optimized using a compact cardiac chamber made of PDMS polymer (SYLGARD 184, DOWSIL) and Petri dishes (Helicon, N-706201). The minimum volume of perfusion of the heart with oxygenated solution was reduced to 10 mL. A syringe pump (Fusion 100 Infusion Pump (Chemix, Stafford, USA)) was used to perfuse the heart with heated Tyrode’s solution during the optical mapping. The contour volume was approximately 50 mL.

The fluorescent signal was recorded with a 64×64 pixels resolution and a sampling frequency of 239 frames per second by the same setting. The heartbeats were removed by addition of BDM (Sigma-Aldrich, B0753-25G) to the solution at a concentration of 10 mM when the heart was stained for fluorescence using Fluo4-AM (Lumiprobe, 1892-500ug) at a concentration of 2.8 ug/ml.

Data were recorded at 37°C using both thermal and electrode stimulation. The duration and amplitude of the electrode and thermal stimulation ranged from 1 ms to 20 ms duration and from 1 V to 6 V. The stimulation frequency ranged from 2 to 4 Hz unless otherwise stated according to the stimulation protocols.The stimulus was set using a generator (2 MHz USB PC Function Generator, PCGU100 (Velleman, Gavere, Belgium)). Platinum electrodes were used

Data processing was undertaken using the ImageJ program and the associated plugins. An ImageJ plugin (time-lapse color-coder) was used to build pseudo-3D images and activation maps. Principal analysis was performed in Wolfram Mathematica 12.

### Raw ECG data analysis

CSV files containing one or two ECG channels and laser trigger pulse channel were generated using PowerGraph with a 10kHz sampling rate. Initial R-peak annotation was performed using *neurokit2*^56^, then the annotation was manually corrected with PhysioZoo interface^57^. Further processing was performed using hand-made python3 scripts with *neurokit2* and standard libraries for data analysis. *scipy*.*signal* and *scipy*.*ndimage* subpackages were used to operate 1D data. Heart rate smooth dependency over time plotted in Fig. 2,3 and others was calculated by applying 1-dimensional Gaussian filter with σ=2s to discrete heart rate data (inverted sizes of RR interval against start time of the R-peak). ECG denoising was performed with PCA (implemented in *scikit-learn* package). Each RR interval was resampled to an average RR length, then PCA was performed and 3 first PCs were used to reconstruct RR interval; next, an interval was resampled back to its original length. During the analysis of pacing data, PCA was performed separately for RR intervals under laser treatment and for control RR intervals. PCA-denoised data were then delineated with *neurokit2* using continuous wavelet transform method to annotate P-peaks.

### Induction of hyperthermia in mice

6-8 week old mice were anesthetized with 1.5% isoflurane and placed on a heating pad in the supine position according to the procedure described in the «Viral vector delivery section». Monitoring of mouse body temperature was performed using a rectal thermometer. To minimize heat loss, mice were covered with a heat-insulating blanket. Then, the temperature of the heating pad was increased from 38°C by 0.5°C every 3 min until the rectal temperature reached above 43°C. ECG was recorded as described in «Electrocardiography of immobilized mice» and ECG data analysis was performed as described in «Raw ECG data analysis section».

### Distant heating systems

Two optical systems were used to stimulate cardiomyocytes *in vitro* and *in vivo*. Both were equipped with a fibre coupled laser diode (LD) 4PN-117 (SemiNex) as a powerful heating laser, providing radiation at a wavelength of 1375 nm with an average power of up to 4.3 W through a multimode fiber with a core diameter of 105 μm and 0.22NA. The LD was mounted onto a TEC-controlled plate “264 TEC HP LaserMount” (Arroyo Instruments, A.I.), which was operated by TEC driver TECSource 5305 (A.I.); current stabilization and control for LD were performed with LD driver LaserSource 4320 (A.I.). For pulse-mode heating, an external pulse trigger was used (see “Electrocardiography of immobilized mice” section).

For cell culture heating an all-fiber system was developed. To illuminate the heating region, we used a PL520 (Thorlabs) laser diode at a wavelength of 520 nm, which was optically conjugated with the tip of a 200-μm 0.22NA FG200UEA (Thorlabs) optical fibre by optical glue. Combining IR and visible radiation was performed using 90:10 splitters TM105R2F2B (Thorlabs) for fibres with a diameter of 105 µm 0.22NA and reduced OH content. For optimal transmission of IR radiation, it was fed into the Signal channel with a 90% power transmission. The splitter’s auxiliary output could be used to monitor the IR power. The main channel was connected to a 2-m-long optical fibre, which supplied IR radiation to the object, with parameters 105 µm/0.22NA (FG105LCA, Thorlabs). Due to low-OH content, the fibre had increased transmission in the IR range, which made it possible to reduce losses by ∼100 times. All light guide elements had connectors, which made it possible to securely connect them to each other. The overall IR power transfer efficiency was 65%. This laser heating system was synchronized with the recording of cell membrane potentials and currents through them using a low-noise amplifier HEKA EPC 10 USB. The end of the fibre was placed on a three-coordinate manipulator (Stoelting). The expansion cone of IR radiation in the solution was ∼9°, which led to an elliptical heating spot of 150×300 µm at a distance of 200 µm from the cells. In experiments on pulsed heating, the IR power reached 2.1 W and the intensity up to 6 kW/cm^2^ (at a maximum control voltage of 2.5 V).

When a heart was stimulated, LD IR laser radiation was free space coupled using an F810SMA-1310 collimator (Thorlabs). This allowed the IR radiation to be combined with emission of a green laser diode at a wavelength of 532 nm CPS532-C2 (Thorlabs) with a power about 1 mW on a DMLP1000 dichroic mirror (Thorlabs) to visualize the heated area. The infrared and visible light were coupled to a 2-m-long FT800EMT (Thorlabs) glass fibre with the core diameter 800 μm and the numerical aperture 0.39NA with a reduced OH content to reduce absorption in the 1350–1500 nm region by a lens with a focal length of 50 mm. To heat the tissue, the end tip of the IR grade light guide was set at a distance of 6 to 8 mm from the surface of the heart, which created a heating spot with a diameter of 4.2 mm to 5.4 mm. The overall efficiency of energy delivery to the heart was ∼70%. In the continuous heating mode, the power varied from 250 mW (controlling voltage is 0.6 V) to 600 mW (1.0 V), which led to heating from 32 °C to 37–43 °C. When heated by pulses, the peak power rose to 2.1 W.

### Inclusion & ethics statement

Our research has been a collaborative effort, including local researchers throughout the entire research process – from study design and implementation to data ownership, intellectual property, and authorship of publications. All collaborators have fulfilled the criteria for authorship required by Nature Portfolio journals and have been included as authors. The research is locally relevant, and this relevance has been determined in collaboration with local partners. Roles and responsibilities were agreed upon amongst collaborators ahead of the research, and capacity-building plans for local researchers were discussed and implemented. Our research was conducted entirely within our country, by a team mostly located in our country. It was not severely restricted or prohibited in our setting. All necessary permissions and approvals were obtained from local stakeholders. The study was approved by our local ethics review committee. All animal procedures were performed in accordance with the ethical standards and local regulations. Our research did not include any health, safety, security, or other risks to researchers. We have taken local and regional research relevant to our study into account in citations. We believe in the importance of recognizing and citing the work of other researchers in our region, and we have made every effort to do so.

## Reporting Summary

Further information on research design is available in the Nature Research Reporting Summary linked to this article.

## Data availability

The main data supporting the results of this study are available within the paper and its Supplementary Information. The entire dataset for the pacing of primary murine cardiomyocytes expressing human TRPV1 can be found in the following repository: https://doi.org/10.5281/zenodo.10402004 (ref. ^28^); and the dataset for ECG recordings of cardiac pacing in mice can be found at https://doi.org/10.5281/zenodo.8346874 (ref. ^29^). Fluorescent images used for the analysis of Ca^2+^ dynamics in cardiomyocytes and studying the localization of hTRPV1 deposited at https://doi.org/10.5281/zenodo.10814759 (ref. ^58^). Plasmid sequences are deposited at https://doi.org/10.57760/sciencedb.16880 (ref. ^59^). Optical mapping data are deposited at https://doi.org/10.5281/zenodo.14774809 (ref. ^60^).

## Code availability

The code for the data analysis is available at https://doi.org/10.5281/zenodo.10813599 (ref. ^61^).

## Acknowledgements

We thank the Group of Redox Biology IBCh RAS, particularly Viktoriya G. Krut’, Liana F. Mukhametshina and Dmitry I. Maltsev for assistance in preparing hTRPV1 (−109 amino acid residues) data. Figure 1A was created in BioRender, Belousov, V. (2025) https://BioRender.com/e10n004.

## Author contributions

A.V.B. conceptualized the work, performed research, developed methodology, verified experimental results, prepared data visualization, and wrote the manuscript; A.M.N. developed methodology, performed research, implemented code, analyzed data, verified experimental results, prepared data visualization, and wrote the manuscript; A.A.L. created all the optical setups; V.S.O. and S.S.S. performed research and wrote the manuscript; E.S.F., S.K.A., and R.M.K. performed research; A.A.Moshch. created viruses; D.J. and R.A.S. performed electrophysiological and calcium imaging experiments, and prepared data visualization; V.D.D., E.A.T., and M.M.S. performed optical mapping experiments, prepared data visualization, and wrote the manuscript; D.Z.B. and A.V.Z. analyzed data, implemented code, and prepared data visualization; G.M.S., E.M.S., and O.V.P. created truncated TRPV1 channel; I.V.K. developed methodology and performed research; V.A.T., K.I.A., A.V.R., and A.B.F. developed methodology; S.V.K. provided medical expertise; T.B., and A.M.Z. conceptualized the work; A.A.Mozh. provided financial support, developed methodology, conducted research, and wrote the manuscript; V.V.B. conceptualized the idea behind the study, and the work, provided financial support, and supervised the study. All co-authors revised and improved the manuscript.

