## Supplementary information for "Thermogenetics for cardiac pacing"

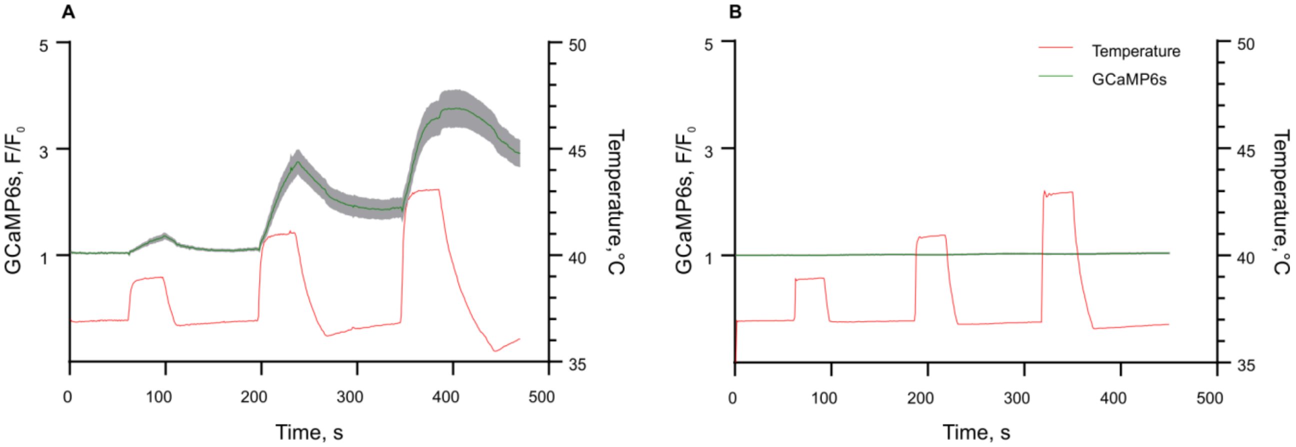


**Supplementary Fig. 1 | Calcium dynamics upon heating in HEK293 cells transfected with hTRPV1(−109aa)** together with GCaMP6s **(a)** and GCaMP6s only **(b).** For **a** 50 cells were measured, and for **b** 54 cells were measured. GCaMP6s fluorescence normalized to the value at the initial time point. The green line indicates median, grey areas show SEM.

**Supplementary Table 1 | Statistics over recordings of successful cardiomyocyte pacing experiments.** Highlighted lines demonstrate the most promising stimulation modes for 2 and 7 Hz pacing. The diversity of experimental conditions is presented in **Supplementary Fig. 2** in a more detailed way. In every case, we tried to find the minimal power required for pacing

| **Frequency, Hz** | **Pulse width, ms** | **Minimal power required for pacing, mW** | **The number of successfully paced cells** | **The number of cells failed to be paced** |
|---|---|---|---|---|
| 2 | 50 | 560 | 5 | 14 |
| | 75 | 600 | 1 | |
| | **100** | **140** | **4** | |
| | 250 | 300 | 3 | |
| 7 | 20 | 330 | 3 | 15 |
| | **30** | **500** | **6** | |
| | 50 | 510 | 1 | |
| **Total:** | | | 23 | 29 |

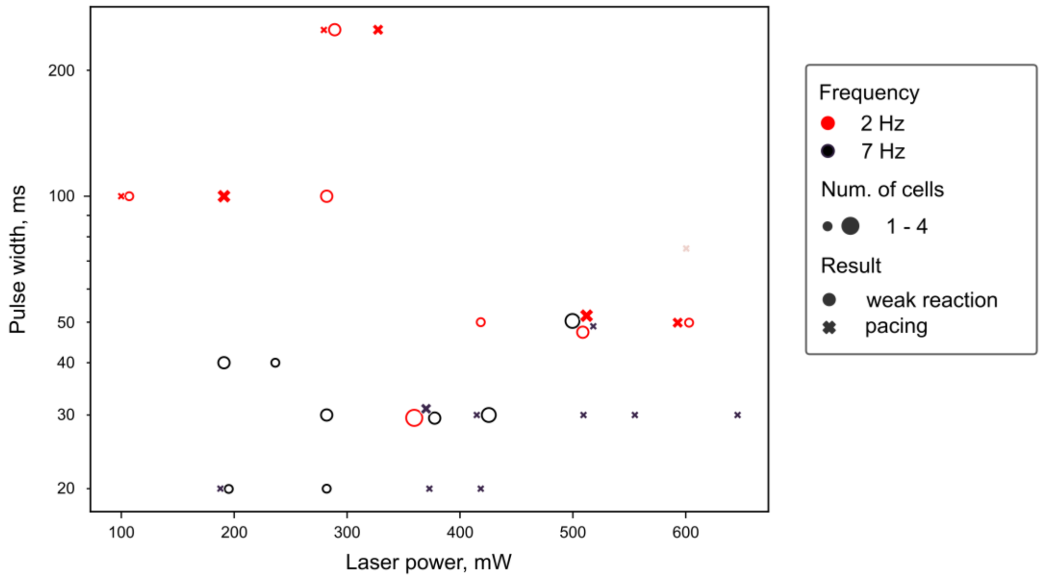


**Supplementary Fig. 2 | Distribution of successful and unsuccessful pacing experiments with cardiomyocytes.** Data is presented depending on stimulation parameters: frequency, pulse width, and laser power.

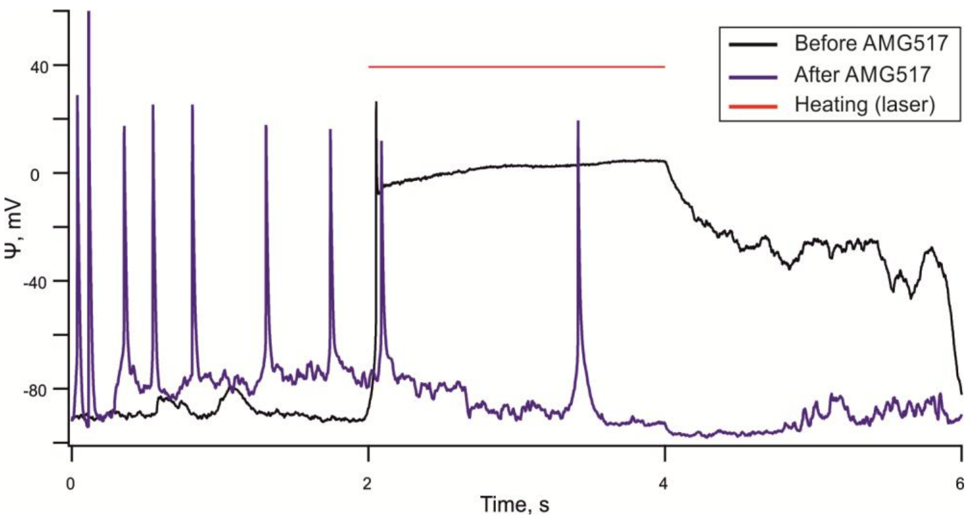


**Supplementary Fig. 3 | Heating of TRPV1-expressing neonatal murine cardiomyocyte with and with AMG517.** The figure shows the same cell before and after application of 1 μM AMG517 solution through a pipette near the cell.

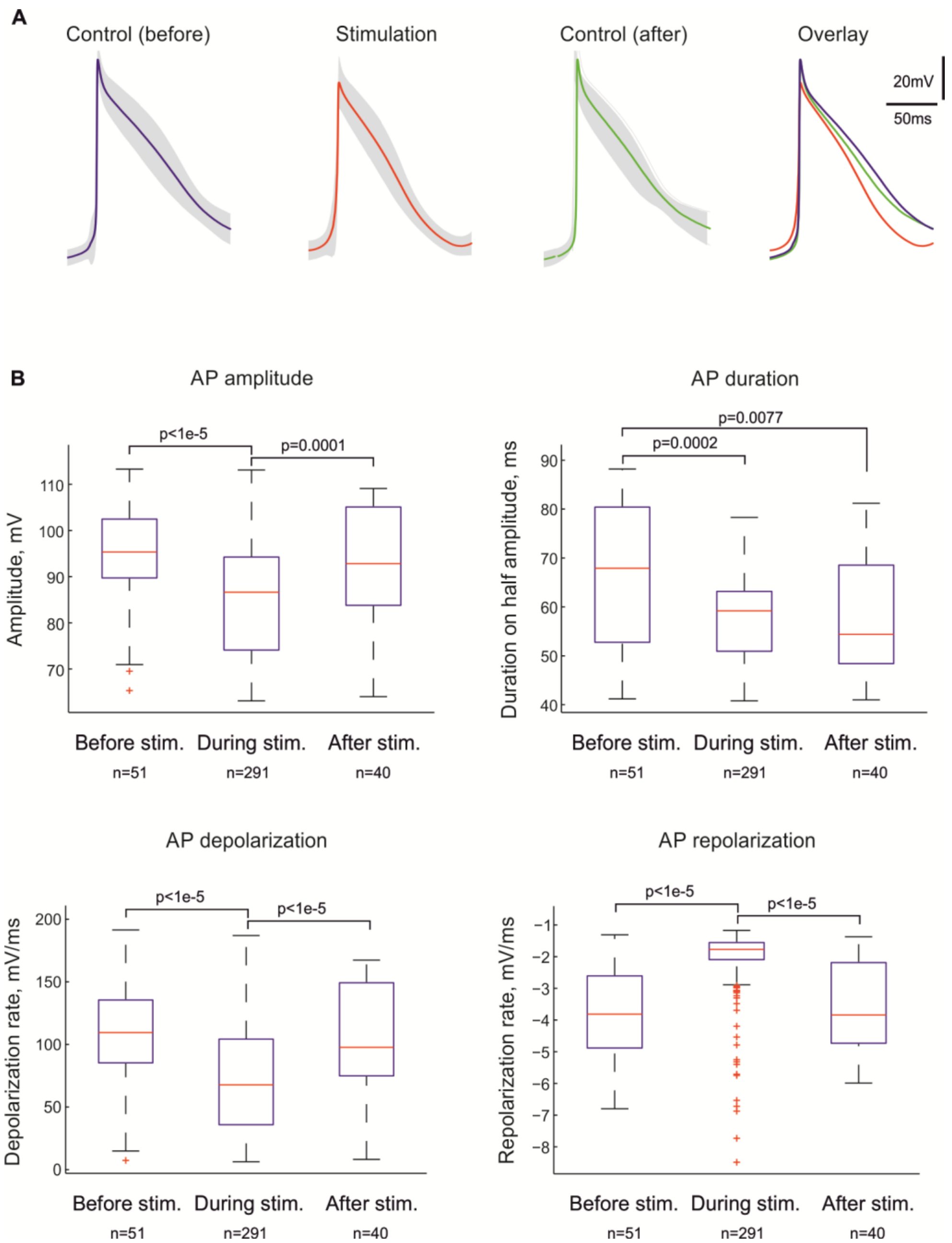


**Supplementary Fig. 4 | Action potential (AP) parameters in TRPV1-expressing murine neonatal cardiomyocytes. a.** Averaged spontaneous and TRPV1-evoked APs (mean±SEM). **B.** Box plots comparing amplitude, depolarization rates, half-width, and repolarization rates of spontaneous APs recorded before and after thermal stimulation with those of TRPV1-evoked APs.

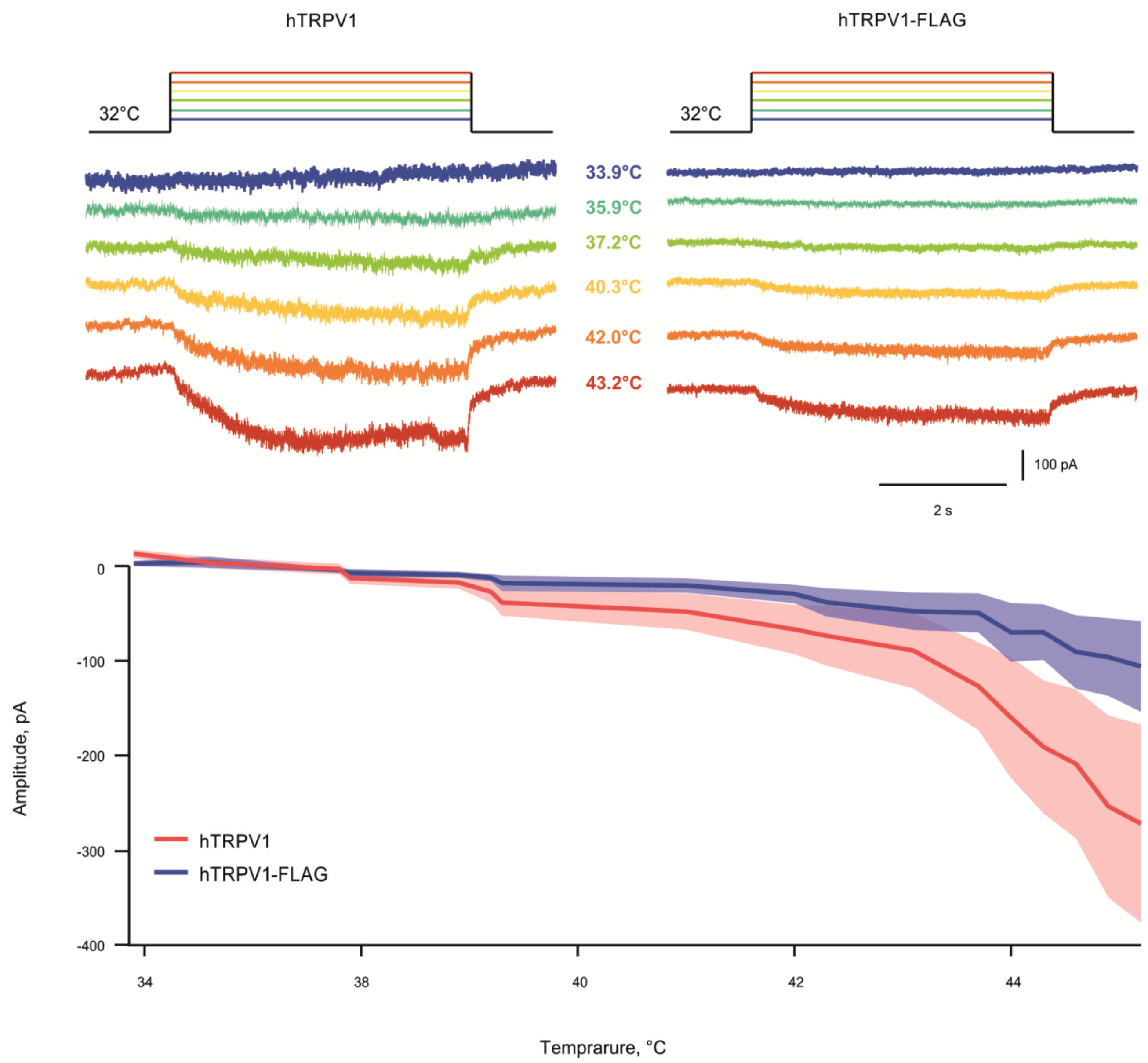


**Supplementary Fig. 5 | Inward currents at different temperatures in HEK293 cells transfected with hTRPV1 and hTRPV1-FLAG.** In the lower graph, lines indicate the median, shaded areas show SEM

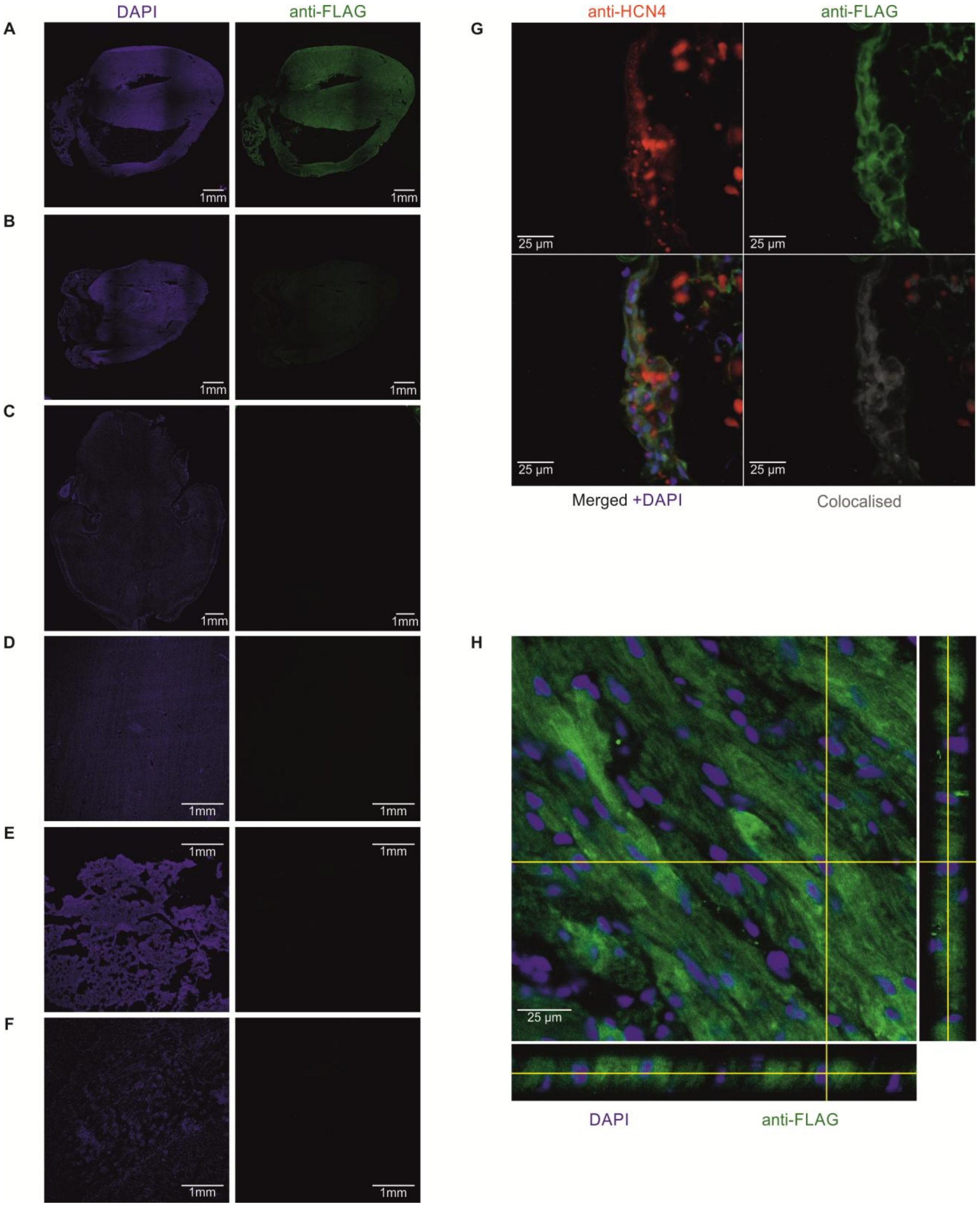


**Supplementary Fig. 6 | Expression and distribution of TRPV1-FLAG in heart and other tissues. a.** Anti-FLAG immunostaining in the heart of a mouse infected with pAAV_cTnT_hTRPV1_3×FLAG vector. **b.** Anti-FLAG immunostaining in the heart of a control mouse. **c-f.** Anti-FLAG immunostaining in the brain **(c),** liver **(d)**, lung **(e),** and skeletal muscle **(f)** of a mouse infected with pAAV_cTnT_hTRPV1_3×FLAG vector. **g.** Co-immunostaining of the sinus node of a mouse infected with pAAV_cTnT_hTRPV1_3×FLAG vector with anti-HCN4 and anti-FLAG antibodies. **h.** Intracellular distribution of anti-TRPV1 immunostaining in the myocardium of a mouse infected with pAAV_cTnT_hTRPV1_3×FLAG vector.

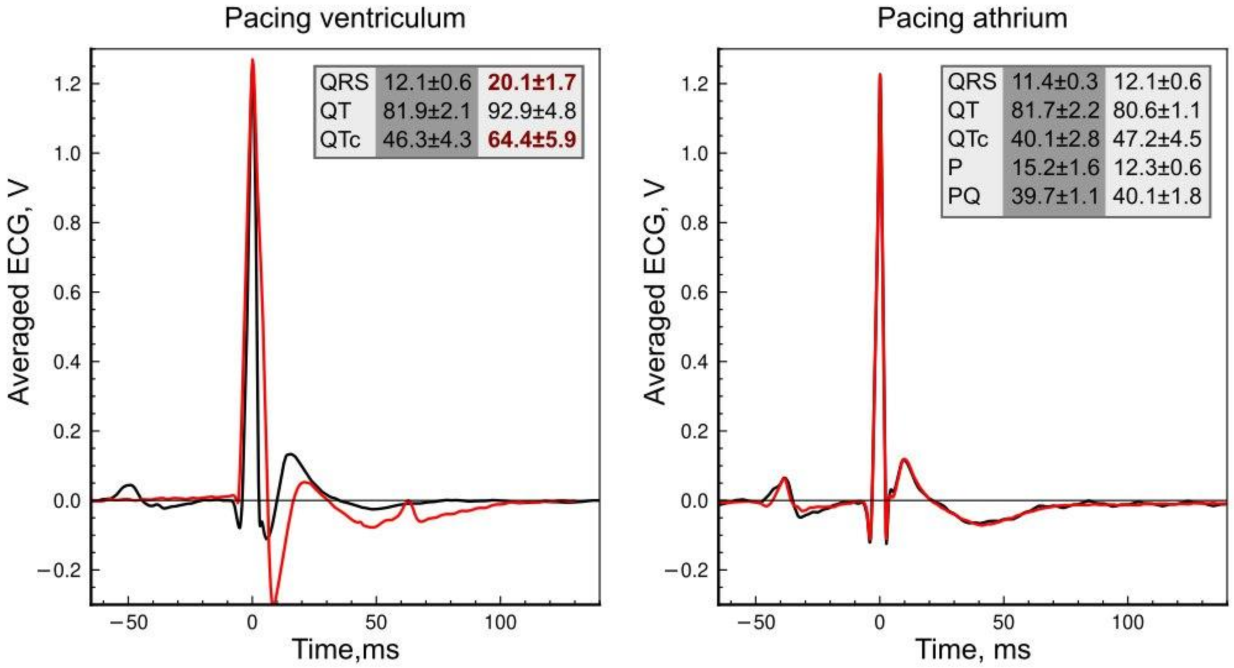


**Supplementary Fig. 7 | Changes in ECG shapes during thermogenetic heart pacing in vivo.** Black curves: ECG shapes before the pacing, red curves: ECG shapes during the pacing. Initial ECG parameters are shown in grey columns, ECG parameters during the pacing are shown in white columns. Significant differences ($p<0.05$) are shown in red.

**Supplementary Table 2 | Statistics over recordings of successful heart pacing experiments.**

| **Heart answer \ Stimulation freq.** | **Number of recordings (Number of animals)** | | | |
|---|---|---|---|---|
| | **2–3 Hz** | **4 Hz** | **5 Hz** | **6 Hz** |
| Locking 1:1 (ventriculum) | 3 (2) | 7 (3) | 4 (3) | 5 (3) |
| Locking 1:1 (atrium) | 2 (1) | 1 (2) | 2 (1) | 4 (1) |
| Incomplete phase locking | 0 (0) | 0 (0) | 3 (2) | 2 (1) |
| Locking 3:2 (atrium) | 0 (0) | 1 (1) | 0 (0) | 1 (1) |
| Locking 4:3 (atrium) | 0 (0) | 0 (0) | 0 (0) | 4 (2) |
| **Total number of animals:** | **4** | | | |
| **Total number of recordings:** | **37** | | | |

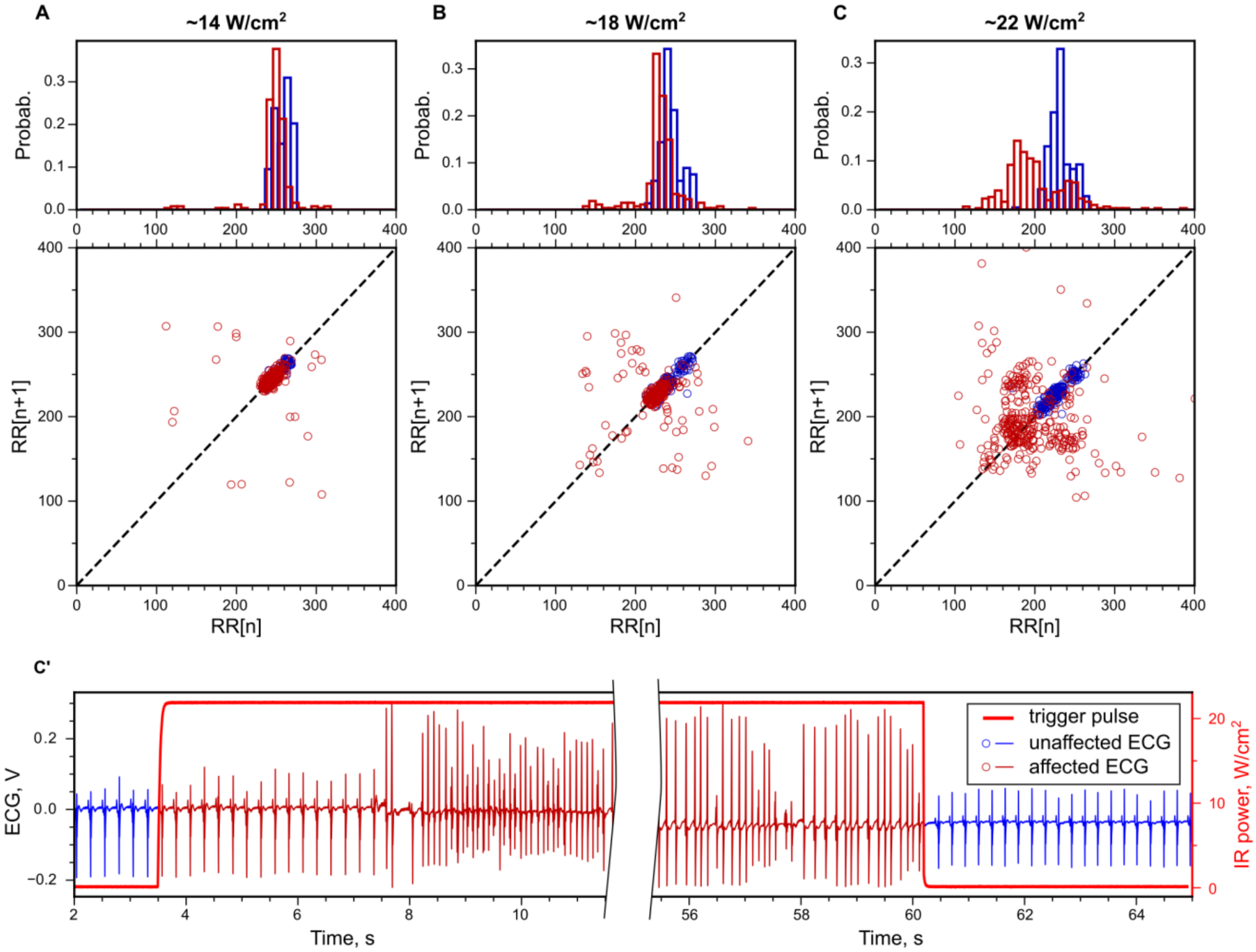


**Supplementary Fig. 8 | Effect of continuous heating of the atrium with three different IR laser intensities on heart rhythm. a–c (top row),** Distribution of RR interval lengths plotted separately for intervals before and after heating (unaffected ECG, blue), and for intervals during heating (affected ECG, red). **a–c (bottom row),** Poincare plots, where each pair of consecutive R-R intervals is a point on a graph. **c',** Denoised ECG corresponding to the 3rd case with the middle part removed.

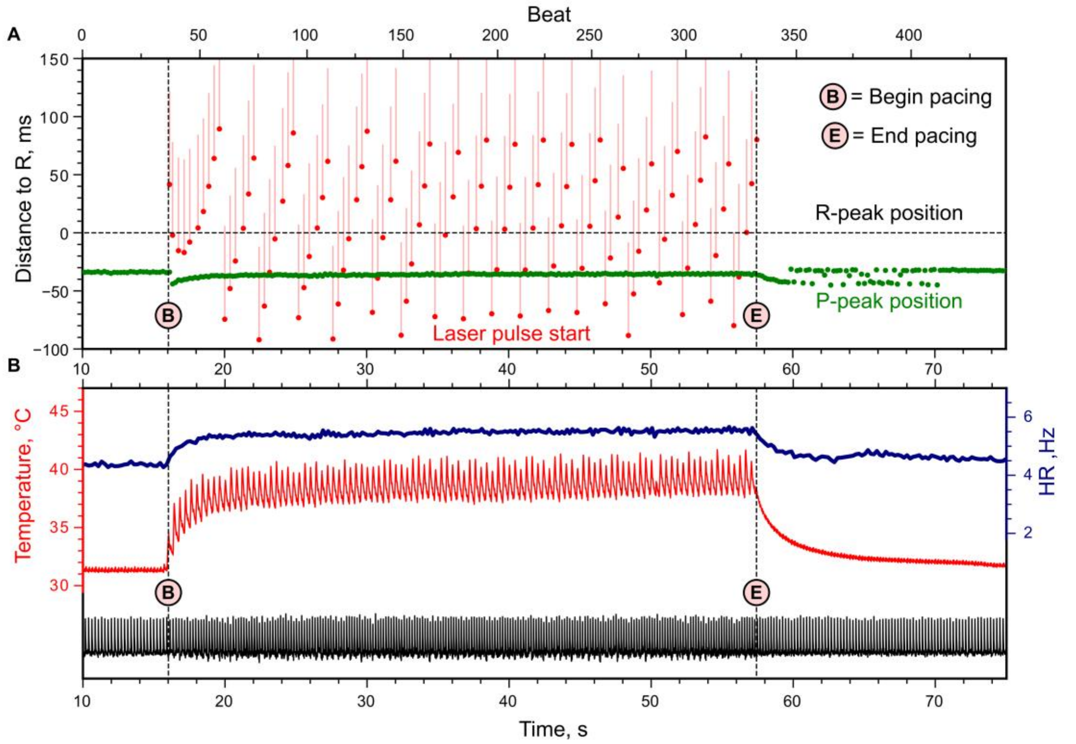


**Supplementary Fig. 9 | Reaction of the heart beating to pulse heating of the atrium in a control mouse.** Pacing was at 2.5 Hz with 80-ms pulses, initial HR ~4Hz. **a.** Position of the P-peak and laser pulse in relation to the R-peak in ms. Laser pulses don't synchronize with the heart beat and are located at different time shifts in relation to R-peak zero line. **b.** Change of ECG signal (black curve), heart rate (blue curve), and temperature (red curve) during the experiment. Temperature was measured using an ultrathin thermocouple probe mounted under the atrium.

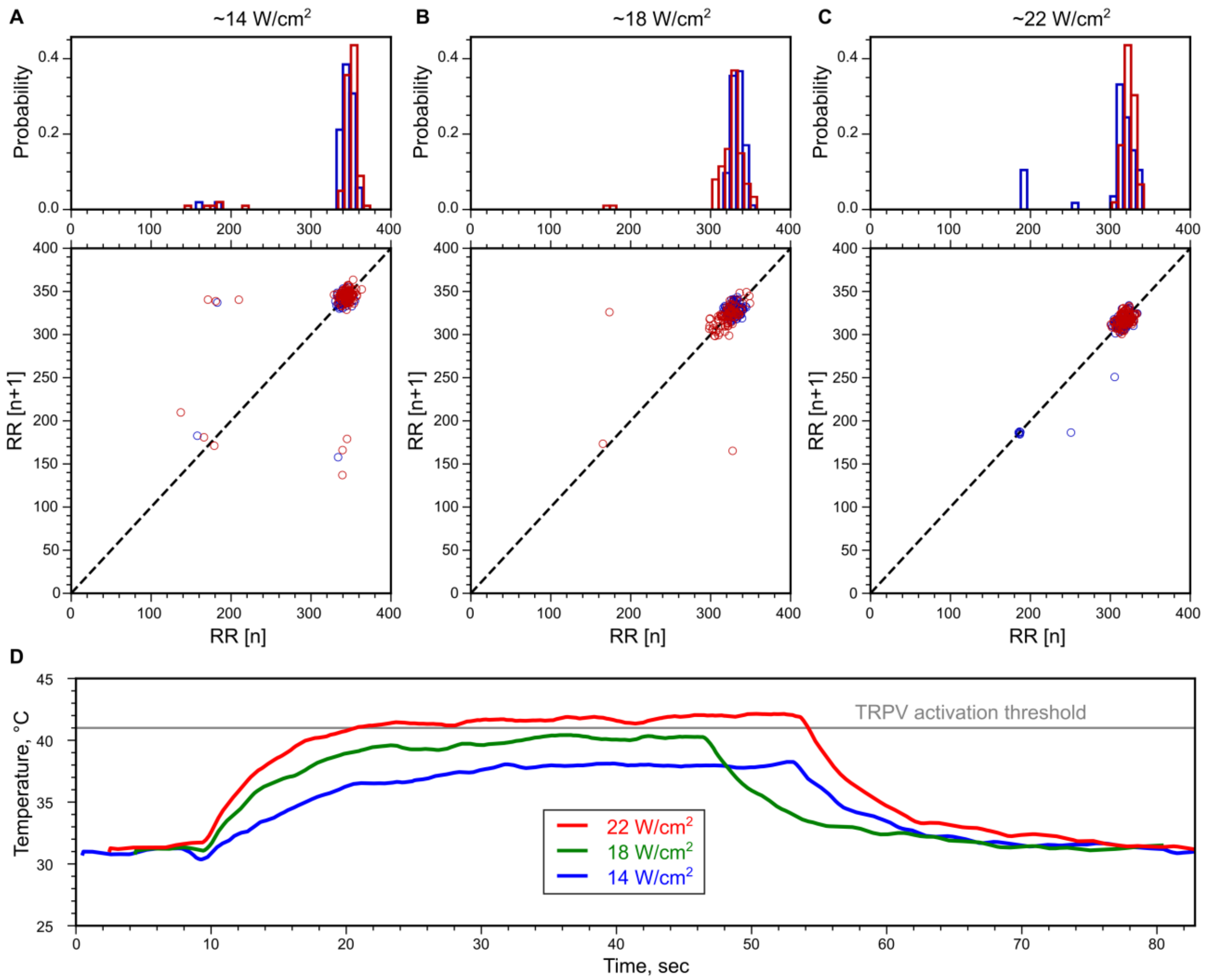


**Supplementary Fig. 10 | Reaction of the heart rate on the constant heat of the heart ventriculum. a–c (top row)**, Distribution of RR interval lengths plotted separately for intervals before and after heating (unaffected ECG, blue), and for intervals during heating (affected ECG, red). **a–c (bottom row)**, Poincare plots, where each pair of consecutive R-R intervals is a point on a graph. **d.** Temperatures during experiments **a–c** were measured using an ultrathin thermocouple probe mounted inside the myocardium.

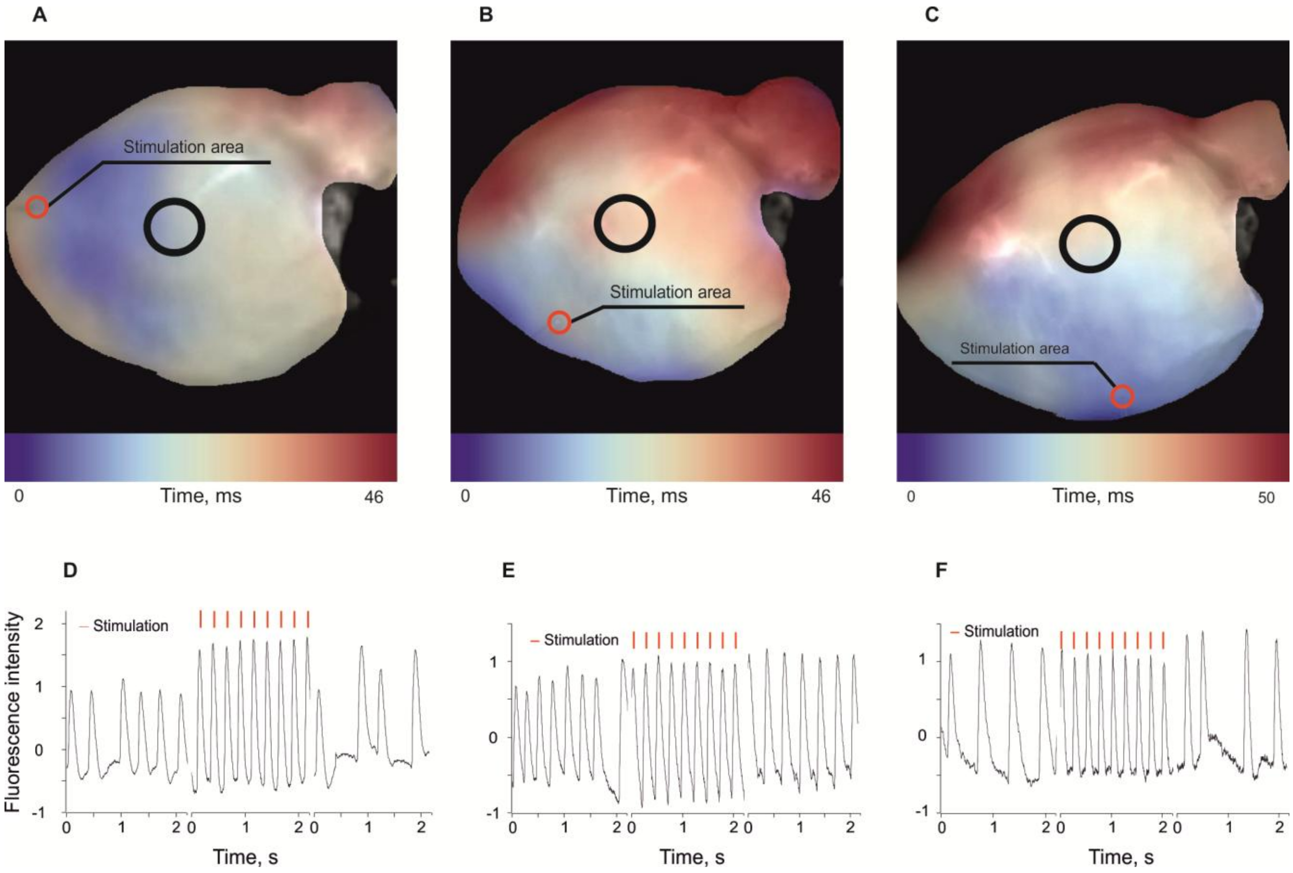


**Supplementary Fig. 11 | Optical mapping analysis of mouse hearts demonstrating the spatial resolution of the method.**

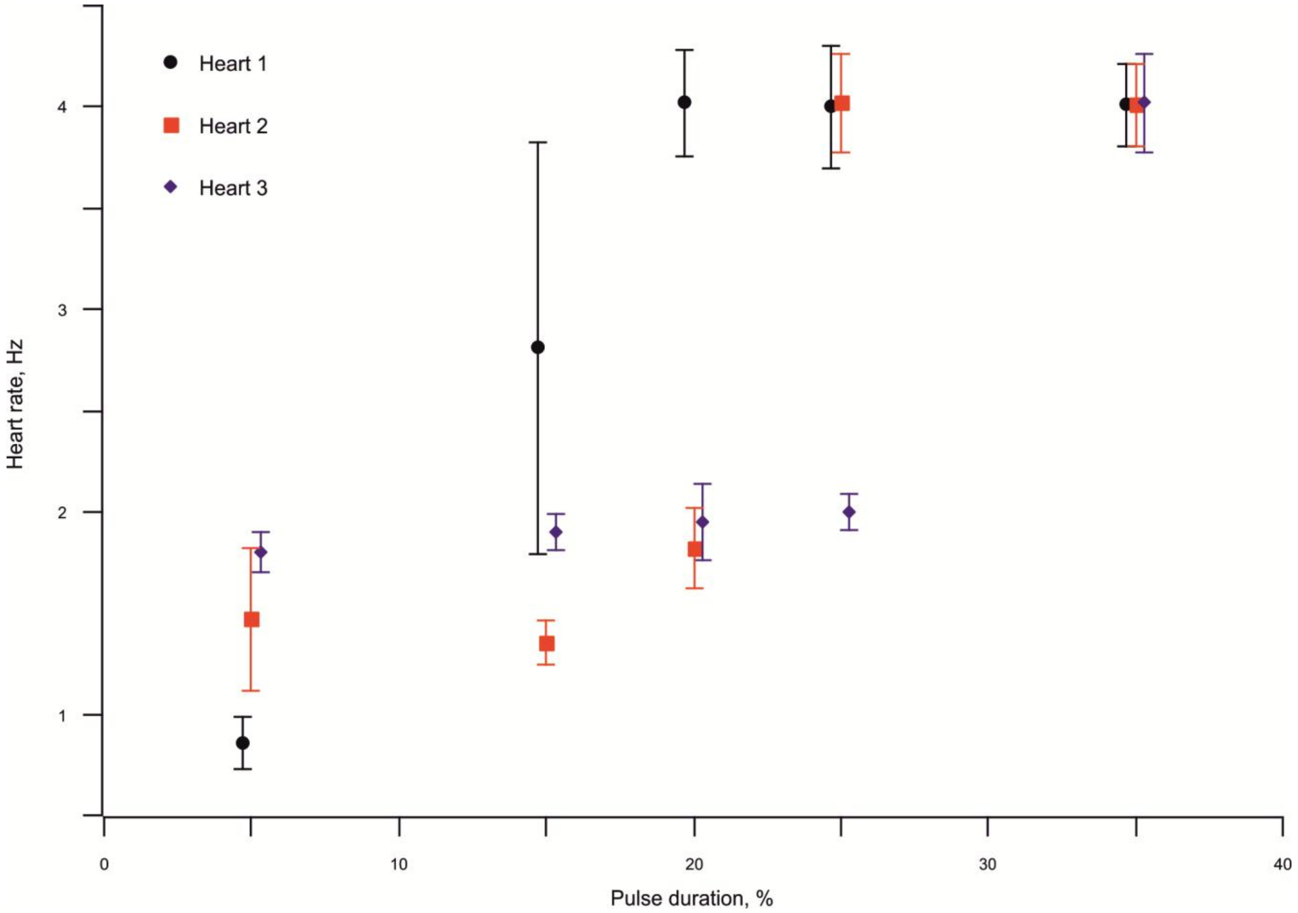


**Supplementary Fig. 12 | Heart apex pacing efficacy depending on the pulse duration.** Stimulation was performed at 4Hz. Pulse duration is shown as a percentage of the interval between two pulses.

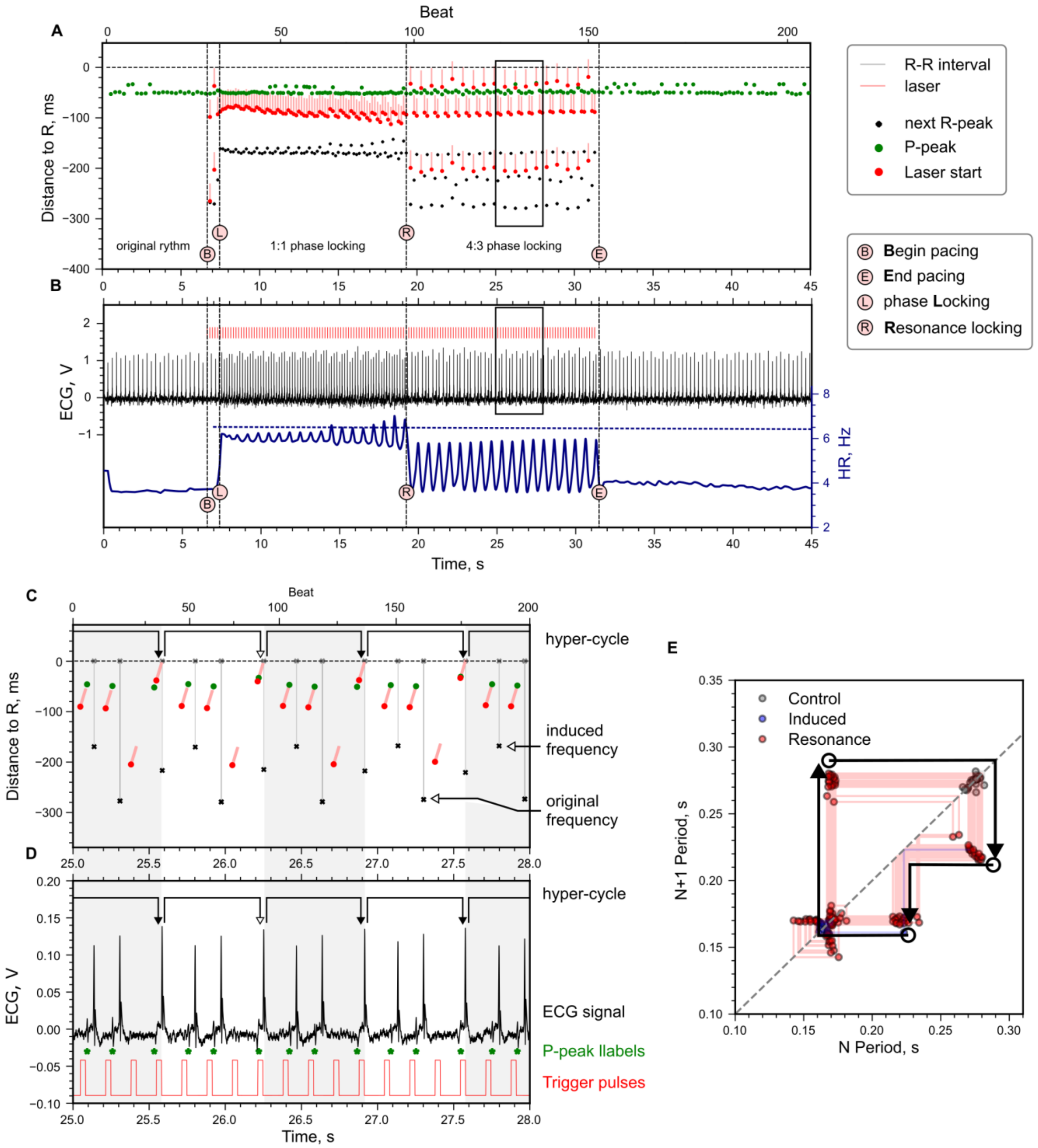


**Supplementary Fig. 13 | Atrial pacing where normal pacing turns to the resonance.** 6 Hz, 33 ms pulses, intrinsic HR ~4 Hz. The 1:1 phase locking turns into 4:4 (each laser pulse corresponds to P and R peaks, but the times between pulses and peaks are different) and then to 4:3. First, every fourth beat occurs earlier than the other three, but every pulse occurs inside a distinct RR interval (locking 4:4). The difference between fourth intervals and others grows, and phase locking changes (event “R”) to 4:3: two laser pulses induce two responses and the next two laser pulses lay within the next R-R **(c)**. The first two pulses end with P-peaks, i.e. they induce atrium contraction, whereas the last two pulses of the cycle lay within 3rd large R-R interval, cumulatively inducing ventricular contraction synchronously with the atrium. The first R-R interval in this 3-point cycle corresponds to the stimulation frequency whereas the last one corresponds to the intrinsic frequency. The heart tries to support both the intrinsic and imposed rhythms if the stimulation is not intense enough. **a,** Position in ms of the P-peak, laser pulse and previous R-peak in relation to the next R-peak. The black frame shows the window zoomed at pane **c**, **b,** Change of complex shape (black curve) and heart rate (blue curve) during the experiment. Red lines demonstrate trigger pulses. **c-d,** Same as **a–b** shown in a larger zoom in a region of 4:3 resonance. Hypercycles consisting of 3 R–R intervals are shown with arrows and a white/light grey background. **e,** Poincare diagram: next R-R interval plotted against previous one. Consequent points are connected by thin lines. The average 3-point hypercycle is shown with bold arrows.

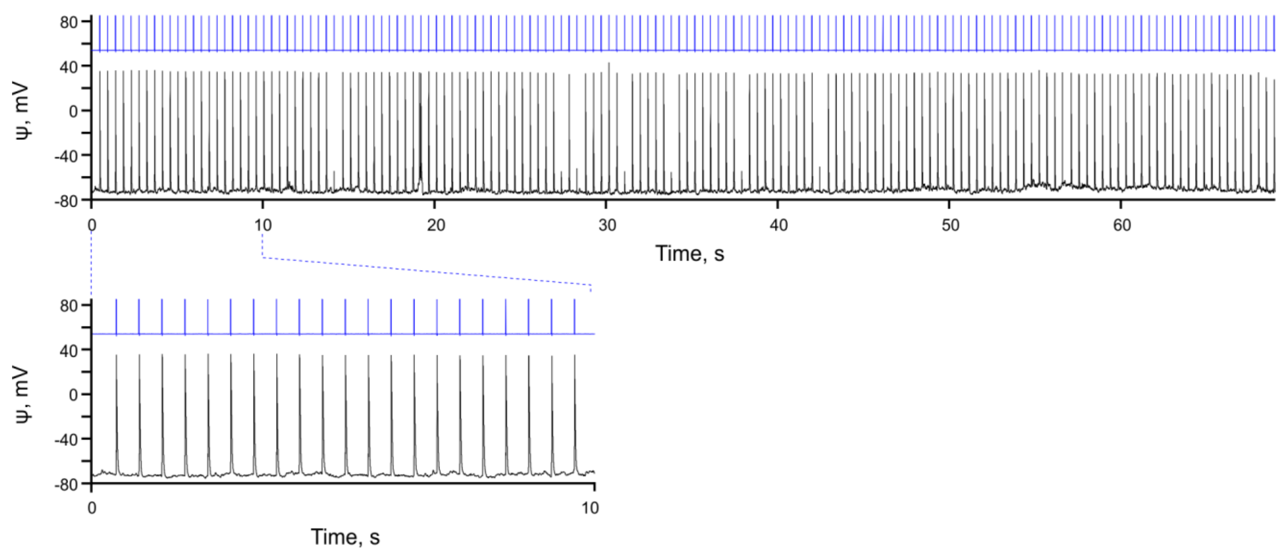


**Supplementary Fig. 14 | Pacing of control mouse neonatal cardiomyocyte with electric pulses.** The membrane potential is shown in black, electric pulses are shown in blue with arbitrary amplitude. Cardiomyocyte recorded in current clamp mode, with 5 ms duration current injections of 30pA.

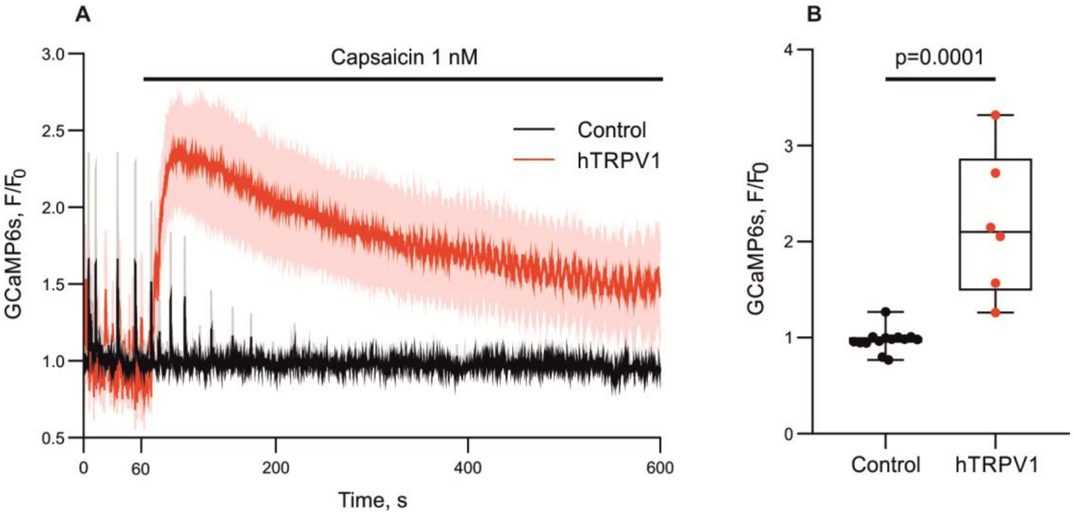


**Supplementary Fig. 15 | Calcium dynamics in a neonatal murine cardiomyocyte transduced with hTRPV1(−109 a.a.) after the addition of capsaicin in a calcium-free medium. a.** Calcium dynamics in hTRPV1(−109 a.a.)-expressing and control cells. **b.** GCaMP6s signal compared 40 s after the addition of capsaicin.

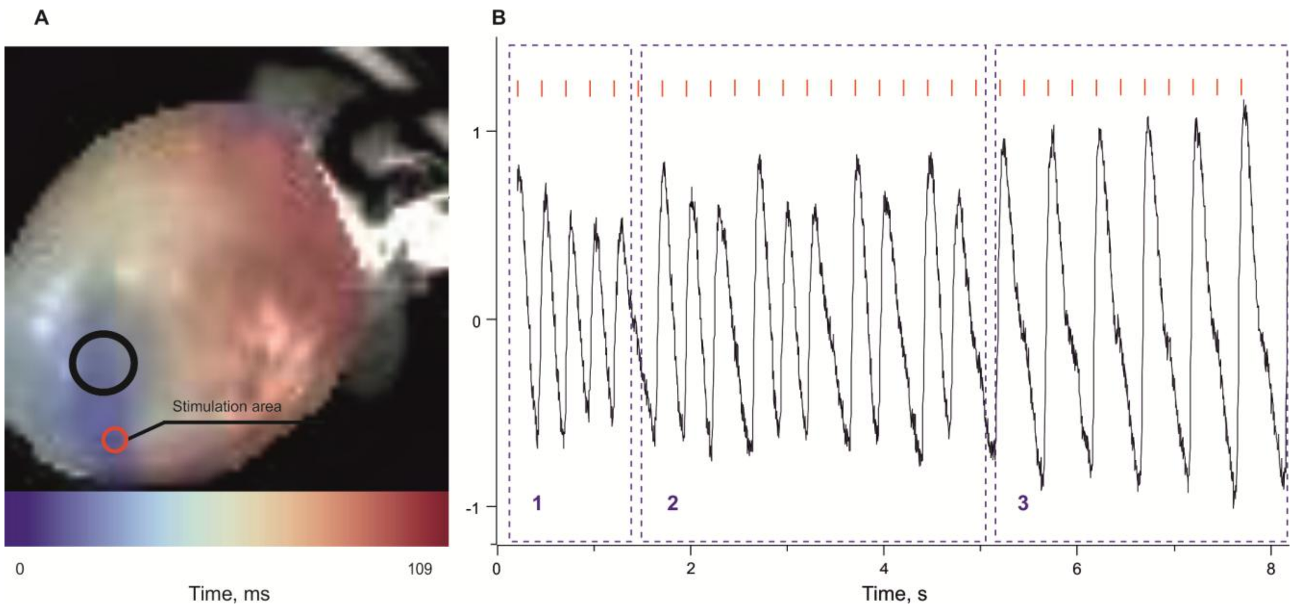


**Supplementary Fig. 16 | The impact of calcium overload induced by 1 mM of caffeine on the efficacy of thermal pacing.** Pacing was performed at 4 Hz. Interval 1 shows the full synchronization (1:1 phase lock). Interval 2 shows 4:3 – 3:2 phase lock. Interval 3 shows 2:1 phase lock.

**Supplementary Table 3 | ECG parameters for gradually heated control (n=3) and TRPV1-FLAG-expressing (n=3) mice.** ECG fragments extracted according to the desired rectal temperature range were processed using PCA filtering and averaged. RR and PQ intervals highlighted in red decreased significantly ($p<0.05$) with temperature.

| Parameter | Control mice | | TRPV1-FLAG mice | |
|---|---|---|---|---|
| | 37–38°C | 41–42°C | 37–38°C | 41–42°C |
| QRS | 11.8 ± 0.5 | 11.1 ± 0.6 | 11.0 ± 0.5 | 11.1 ± 0.4 |
| QT | 45.6 ± 2.7 | 43.5 ± 1.0 | 42.2 ± 6.2 | 37.7 ± 3.8 |
| QTc | 44.4 ± 1.7 | 45.2 ± 0.5 | 39.7 ± 4.6 | 38.9 ± 3.3 |
| P | 10.2 ± 0.8 | 9.4 ± 0.8 | 8.1 ± 2.5 | 8.2 ± 0.7 |
| PQ | 36.0 ± 1.2 | 31.3 ± 1.0 | 35.4 ± 1.0 | 31.1 ± 1.5 |
| RR | 105.3 ± 7.7 | 92.6 ± 3.1 | 111.4 ± 7.3 | 93.6 ± 4.7 |

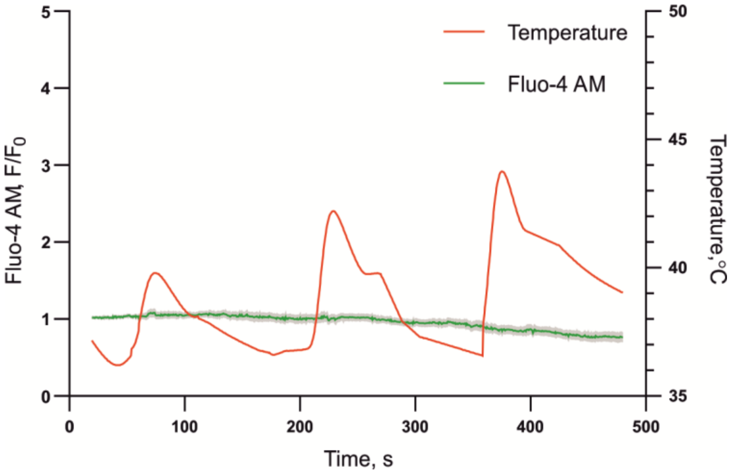


**Supplementary Fig. 17 | Calcium dynamics upon heating in non-myocardial cells isolated from neonatal mouse hearts.**

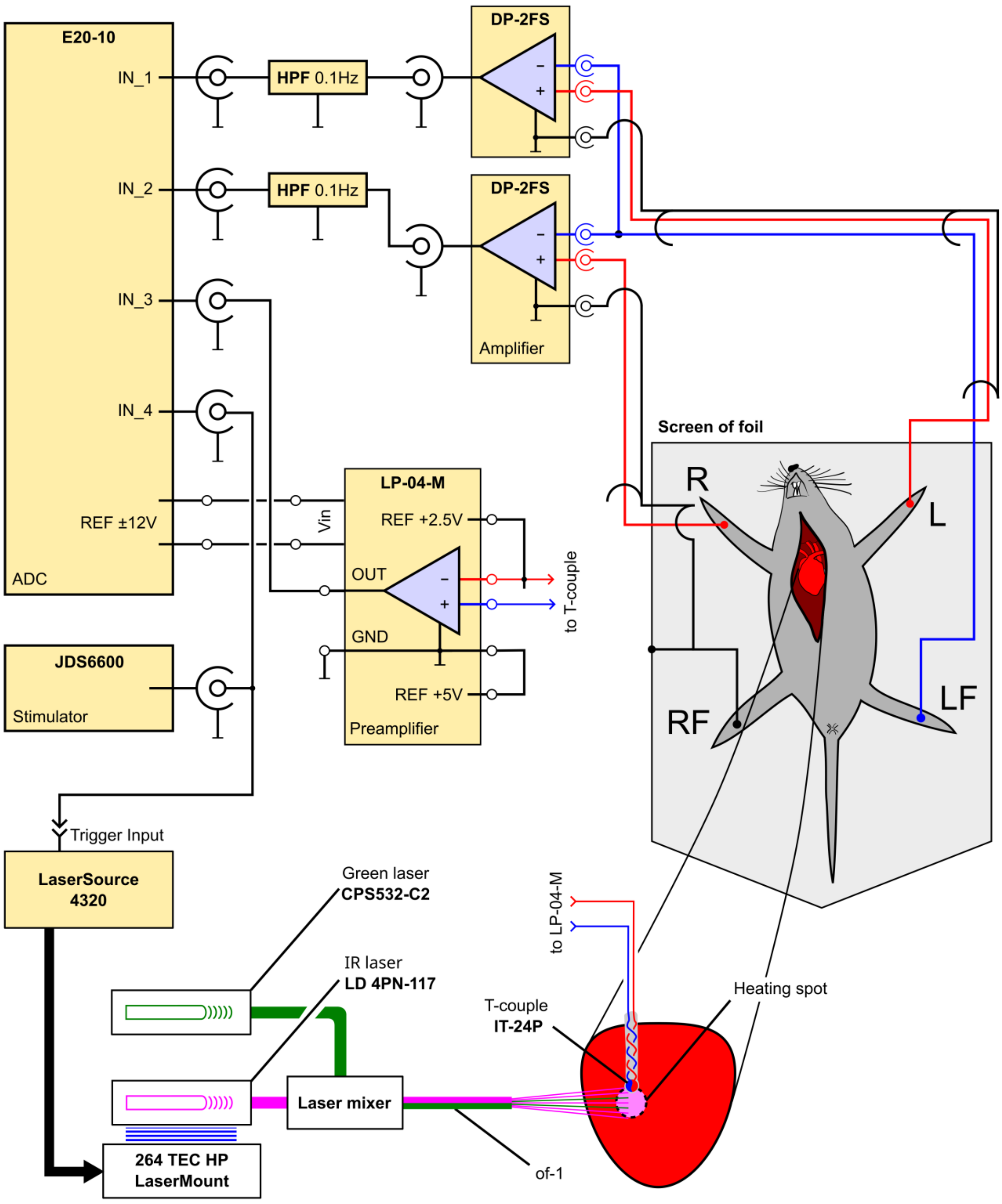


**Supplementary Fig. 18 | Interconnected electrical components allowing precise monitoring of heart rhythm and IR laser pulses.** Signals from ECG electrodes at four leads (**R**, **L**, **RF**, and **LF**) were processed with two **DP-2FS** band differential amplifiers (shown). The two resulting amplified single-ended signals (**R**–**LF** and **L**–**LF**) were processed with a simple high-pass filter (**HPF**, shown) and fed to the 1st and 2nd inputs of the **E20-10** ADC (shown), which was connected to a PC by a USB interface. The **JDS6600** stimulator (shown) generated trigger pulses and operated laser diode driver **4320** (shown); these pulses were recorded by the 4th input of the ADC. The driver fed IR laser diode **4PN-117** (shown), whose light was mixed with a visible green light from **CPS532-C2** source (shown) by means of dichroic mirror (**Laser mixer**) and focused into the optical fibre (**of-1**). The fibre was mounted on a holder in close proximity to the heart, allowing the **heating spot** (shown) to be rather small. The ultrasmall thermocouple **IT-24P** (shown) was inserted into the myocardium at the location of the heating spot; its signal was processed by an **LP-04-M** preamplifier (shown) and fed into the 3rd input of the ADC.
